# Reversed Signaling Flow of a Bacterial Pseudokinase

**DOI:** 10.1101/812529

**Authors:** Kimberly A. Kowallis, Elayna M. Silfani, Amanda P. Kasumu, Grace Rong, Victor So, W. Seth Childers

**Affiliations:** Department of Chemistry, University of Pittsburgh, Pittsburgh, PA 15206, USA

**Author notes:** Corresponding author; W. Seth Childers.

## Abstract

Bacteria respond to environmental and cellular cues both through isolated signaling events between one sensor histidine kinase and its response regulator, and through more interconnected arrays. *Caulobacter crescentus* achieves asymmetric division through a network of histidine kinases, and here we interrogate a novel DivL pseudokinase reverse signaling mechanism that enables productive cross-talk across the network. A leucine zipper fusion method was used to synthetically stimulate reverse signaling between the sensor and kinase domains and directly test if reverse signaling could modulate the signaling network *in vivo*. Stimulation of sensor-kinase helix conformational changes resulted in changes in *C. crescentus* motility and DivL accumulation at the cell poles. The repurposed roles of the sensor domain in these processes were evaluated. We demonstrate that a domain of unknown function that binds to two scaffolding proteins, and two conserved signaling domains are employed as modulators of an active kinase. We propose that reversed signaling may be widely used across signaling enzymes.

## Introduction

Bacterial signaling systems are often arranged as parallel signaling arrays that include a set of histidine kinases (HK) that strictly regulate the phosphorylation of their corresponding response regulator (RR) (Figure 1A) (1). These signaling arrays offer orthogonal connectivity of single signals to single output responses yet have limited abilities to integrate multiple signals. In contrast, productive cross-talk within multi-kinase networks (MKNs) appears to integrate many signals to regulate developmental processes such as sporulation (2), biofilm formation (3), quorum sensing (4), asymmetric cell division (5) and multi-cellular fruiting bodies (6). Studies of these developmental networks point towards kinase-kinase interactions playing crucial roles in signal integration (5, 7–12). Despite the potential importance towards developmental regulation, the mechanisms of kinase-kinase regulation are poorly understood.

**Figure 1:**
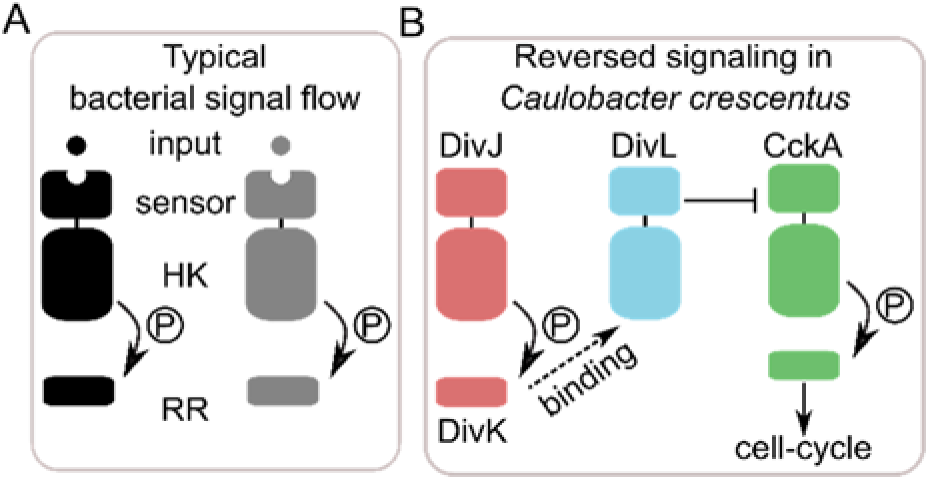
*Caulobacter crescentus* uses the pseudokinase DivL to reverse canonical signaling. A cartoon comparing canonical forward histidine kinase signaling to the DivL pathway in *C. crescentus*. (A) In typical bacterial signal flow, orthogonal histidine kinases receive a signal with the sensor domain, triggering a conformational change and auto-phosphorylation of the HK domain. The phosphate group is transferred to a corresponding response regulator, which elicits a distinct cellular response. (B) In the DivL pathway, the typical histidine kinase DivJ phosphorylates its response regulator. DivL then recognizes phosphorylated DivK through a specific binding event and allosterically modulates the typical hybrid histidine kinase CckA, which prompts a cell-cycle regulating phosphorelay.

A well-studied bacterial multi-kinase network regulates asymmetric cell division in *Caulobacter crescentus* by regulating genes associated with division, replication and motility (Figure 1B) (13–16). This system involves at least four HKs, three RRs, and a phosphotransfer protein that regulate the phosphorylation of the cell-cycle master regulator CtrA (17). Signaling cross-talk amongst four kinases is achieved by an unorthodox bacterial pseudokinase DivL (Figure 1). While DivL is essential to cell survival, the minimal functional DivL lacks most of the catalytic domain (18), and promotes cross-talk without requiring DivL phosphorylation (19–21). Biochemical studies have demonstrated that DivL’s kinase domain binds its cognate response regulator DivK~P (5, 19), but DivL does not exhibit kinase or phosphatase functions towards DivK (5).

Studies of pseudoenzymes in eukaryotic systems have highlighted that while pseudokinases have lost critical catalytic functions, they can impact biochemical systems by functioning as allosteric modulators of active kinases, competitors for substrate binding, subcellular localization anchors and signal integrators (22). Past studies have indicated that DivL shares many features with its eukaryotic counterparts including DivL’s ability to modulate CckA activity in response to DivK~P binding (8, 12). Moreover, DivL plays a role in anchoring CckA at the cell pole (8), which is critical for asymmetric regulation at each cell pole (23). Here we aim to examine the mechanism of the DivL pseudokinase further in order to extract more general design principles of pseudoenzymes. Past studies have suggested that DivL’s kinase domain has been repurposed as a phosphospecific response regulator sensory domain (24). Critically, DivL’s binding to DivK leads to repression of the CckA-CtrA signaling pathway through a mechanism other than direct DivL-CckA phosphotransfer (5, 12).

Sensory domain interactions between CckA and DivL coordinate DivL’s regulation of CckA. Construction of a chimeric CckA kinase demonstrated that CckA’s sensory domains were sufficient for DivL-dependent stimulation in *C. crescentus* (21). In addition, *in vitro* studies of the DivL-CckA complex upon liposomes has shown that DivL and CckA’s sensory domains are needed for DivL to regulate CckA function (25). From these data, we hypothesize that DivL has also repurposed its sensor domain as a regulator of CckA signaling. Together this leads to a reversed signaling model in which DivK~P binding to the pseudokinase domain activates the sensory domain as an “effector” of CckA kinase activity (Figure 1B) (26). A lack of experimental strategies to directly test if reverse signaling occurs and is critical within a cellular context motivated the methodologies described in this study.

When a signal binds to the sensory domain of a typical HK, the sensory domain transmits signals through a coiled-coil linker that allosterically alters the downstream kinase helical bundle conformation (27, 28). A set of conserved D-(A/I/V)-(T/S)-E residues at the junction of the per-ARNT-sim (PAS) sensory domain and this coiled-coil linker forms a network of hydrogen bonds with the PAS domain fold (29, 30). This junction critically couples the conformational change that occurs in the PAS domain upon signal detection to the conformational re-arrangement that allows HK autophosphorylation (31–33). Thus, models for allosteric conformational changes from the PAS sensory domain to the HK domain are now well established. However, it has not yet been shown that this set of allosteric conformational changes can work in reverse amongst this histidine kinase fold family.

Protein engineering strategies provide an approach to allosterically stimulate proteins or disrupt conformational changes and test the reverse signaling model. One approach involves replacing the sensory domain with a leucine zipper with a strong bias to one dimerization interface from the *Saccharomyces cerevisiae* transcriptional activator GCN4 (34). This approach was applied to a chemotaxis receptor Tar (35) and the *Agrobacterium tumefacians* histidine kinases VirA (36–38) and *Staphylococcus aureus* histidine kinase AgrC (39, 40) and demonstrated that covalent attachment of the leucine zipper can stabilize and "lock" the weaker sensory helix coiled-coils into a distinct activity state. We hypothesized that DivL utilizes its helical sensory-HK linker and conserved PAS domain signal transmission motif in a similar but reverse manner. In this case, DivK~P binds the catalytically dead kinase domain which induces a bending or twisting in the sensory helix. This conformational change is transferred to the repurposed sensor through the rigid signal transmission motif, resulting in a change in the sensor’s effect on CckA. Here we apply this leucine zipper fusion strategy at the C-terminus of the DivL sensory domain and use signal transmission motif mutations strategy to interrogate the reverse signaling model (Figure 1) using *in vivo* assays. We then reveal roles of the sensory subdomains as repurposed cell-cycle and localization effectors.

## Results

### DivL must retain its sensory helix to impact upon swarm size regulation

Previous studies have demonstrated that a large portion of the DivL histidine kinase domain (HK), residues 566-769 (18), is not essential for cell viability (18, 20). The remaining minimal functional DivL maintains the multi-PAS sensory domain, the sensory helix, and about one-half of the DivK~P binding site (Figure 2A). It is possible that the DivL(1-565) variant maintains a sufficient interface to interact with DivK~P (5), so we designed two C-terminal truncations to remove the entire HK and the sensory helix. DivL(1-537), hereafter called DivL-537, lacks the entire HK and retains the majority of the sensor helix (Figure 2A). DivL(1-518), hereafter called DivL-518, lacks the sensor helix and only retains the multi-PAS sensory domain (Figure 2A). To evaluate the function of these constructs, we employed a *C. crescentus* DivL overexpression swarm assay that is sensitive to CtrA-regulated physiology that includes motility, division and replication defects (41). We initially confirmed that induction resulted in over-expression of each variant via Western blot analysis (Figure S1). Compared to a strain replicating an empty vector, overexpression of wild-type DivL reduced swarm size to 47 ± 3% (Figure 2B). In contrast, overexpression of DivL-518 resulted in a loss of the small swarm size phenotype characteristic of DivL overexpression (Figure 2B). These results indicate that the sensor helix is critical for DivL’s overexpression impact on cell-cycle progression and, critically, the DivK~P binding site is non-essential for this function.

**Figure 2:**
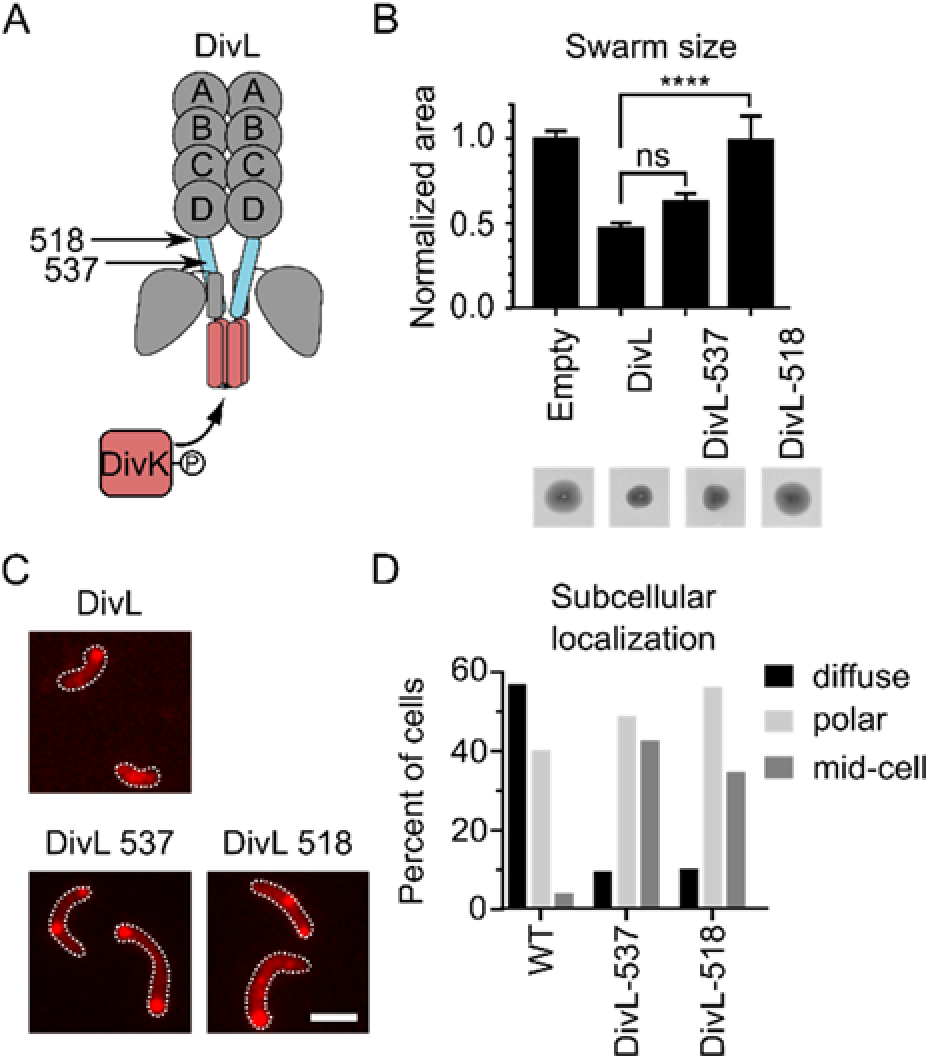
DivL transmits signal between the sensor and HK domains. (A) Model of DivL’s domain architecture with the linker that connects the sensor and the HK shaded blue and the DivK binding site shaded red. (B) Quantification of the motility assay of *C. crescentus* strains expressing DivL-M2 DivK binding site truncations. Representative swarm images are show beneath the corresponding bar. *C. crescentus* were stabbed into 0.3% PYE agar supplemented with 0.3% xylose and incubated at 28 ºC for 3 days. Error bars represent SD (N=3). Significance was determined with a Dunnett’s multiple comparisons test by comparing the mean area of each strain to the mean area of DivL-WT (ns: P>0.05, *: P ≤ 0.05, **: P ≤ 0.01, ***: P ≤ 0.001). (C) Fluorescence microscopy to visualize the subcellular localization of DivL-mCherry DivK binding domain truncations expressed in *C. crescentus*. The variants were induced from the chromosomal xylose promoter in M2G supplemented with 0.03% xylose for 4 hours. Scale bar denotes 2 µm. (D) Quantification of the number of cells with diffuse, polar, or mid-cell localization of cells. N>101 cells for all samples.

### DivL’s sensor helix is not required for foci accumulation at the poles

Previous studies have demonstrated the importance of subcellular localization to DivL’s role in asymmetric cell division in *Caulobacter crescentus* (19, 21, 42). As previously reported, wild-type DivL can be diffuse or accumulate at one or both cell (Figure 2C-D). In comparison, we observed that DivL-537 and DivL-518 exhibited similar levels of foci at the cell pole, however contained a significant increase in mid-cell localized foci (Figure 2C-D). These results indicate that the sensor has the ability to form foci at the cell pole, and that the HK domain plays in regulating subcellular positioning. This observation is consistent with earlier work, that has shown that the C-terminal residues of DivL are involved in DivL’s new cell pole localization (43) (8, 44). Moreover, we also observed that the linker composition between the HK domain and the fluorescent protein fusion impacted cell pole binding (Figure S2). When we observed monopolar DivL variants they were frequently observed at the old cell pole, however did not substantially impact fitness when presented as a sole copy (Figure S2). Collectively, these results indicate that the sensory domain is sufficient for cell pole accumulation, and that the C-terminal residues of DivL strongly influence DivL’s subcellular positioning.

### DivL utilizes a conserved PAS domain signal transmission motif

Because truncating the sensor helix had an effect in the swarm assay, we analyzed this region for conservation amongst DivL alpha-proteobacterial homologs (Figure S3A-B). By mapping the conserved residues onto the DivL HK crystal structure (5), we found high conservation at coiled-coil interface. This suggests that the helix may serve a role in signal transduction between the HK and sensory domains. The alignment also revealed a conserved PAS signaling transmission motif, at residues 516-519 that is critical for transmission from the PAS sensory domain to the HK domain (Figure 3A-B) (29, 45). To test if signal transmission between the sensor and HK domains is critical for DivL’s function, we incorporated mutations that disrupt functionality of the PAS signal transmission motif and examined their impact upon DivL overexpression mediated swarm size reduction. Overexpression of wild-type DivL, confirmed by Western blot analysis (Figure S1), reduced swarm sizes to 55 ± 11% level of the empty vector control (Figure 3C, S4). In contrast overexpression of the DivL D516A and T518V mutants partially recovered the wild-type swarm phenotype with milder reductions in swarm size of 77 ± 3% and 71 ± 5%, respectively. These data indicate that a functional DivL PAS-D signal transmission motif plays a role in the reduction of swarm size upon DivL overexpression. In contrast mutations of the signal transmission motif had only a mild impact upon DivL’s subcellular localization (Figure S5).

**Figure 3:**
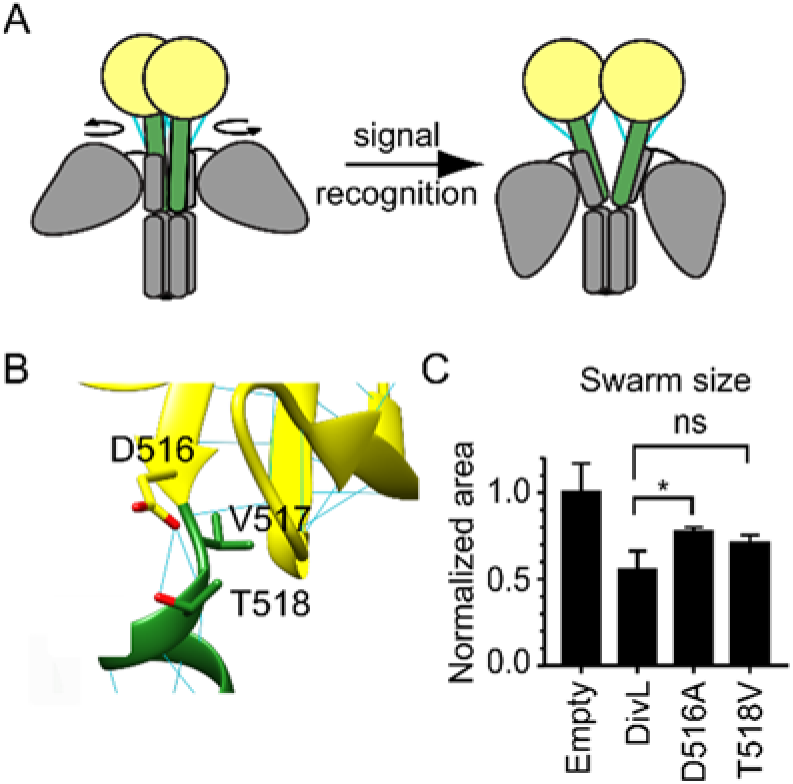
The conserved signal transmission motif is critical for DivL activity. (A) Cartoon of the conformational change that occurs at the signal transmission motif of sensor kinases. The signal transmission motif between the sensor (yellow) and the coiled-coil linker (green) contains residues that form several hydrogen bonds (blue) and serve as a conformational switch. (B) Homology model of DivL compared to YF1 (PDB ID: 4GCZ-A) (33) (C)) Quantification of the motility assay of *C. crescentus* strains expressing DivL-M2 signal transmission motif mutations. *C. crescentus* were stabbed into 0.3% PYE agar supplemented with 0.3% xylose and incubated at 28 ºC for 3 days. Error bars represent SD (N=3). Significance was determined with a Dunnett’s multiple comparisons test by comparing the mean area of each strain to the mean area of DivL-WT (ns: P>0.05, *: P ≤ 0.05, **: P ≤ 0.01, ***: P ≤ 0.001).

### Design of leucine zipper fusions strategy to synthetically trigger reversed signaling

To further interrogate the DivL reverse signaling model, we needed a strategy to synthetically activate the sensor domain and monitor downstream signaling effects. Previous studies have applied fusion of the rigid leucine zipper to the coiled-coil sensory helix to "lock" the kinase in an active, neutral, or inactive conformation (Figure 4A) (36–40). We hypothesized that the leucine zipper could similarly lock DivL’s multi-PAS sensor effector domain into a series of conformations with varying effects upon CtrA-mediated phenotypes and subcellular localization (Figure 4A). To optimize the dynamic range of the response to the leucine zipper fusion sites, we chose DivL-537 as the core construct due to its moderate activity in the swarm assay (Figure 2B) and its ability to form foci (Figure 2C-D). The fusion site of the rigid leucine zipper to the sensory helix of an HK determines the position of its coiled-coil interface, as the addition or removal of one residue results in a rotation of 102°. We thus designed a series of DivL-leucine zipper fusions (DivL-LZ), denoted by their DivL fusion site (Figure 4B).

**Figure 4:**
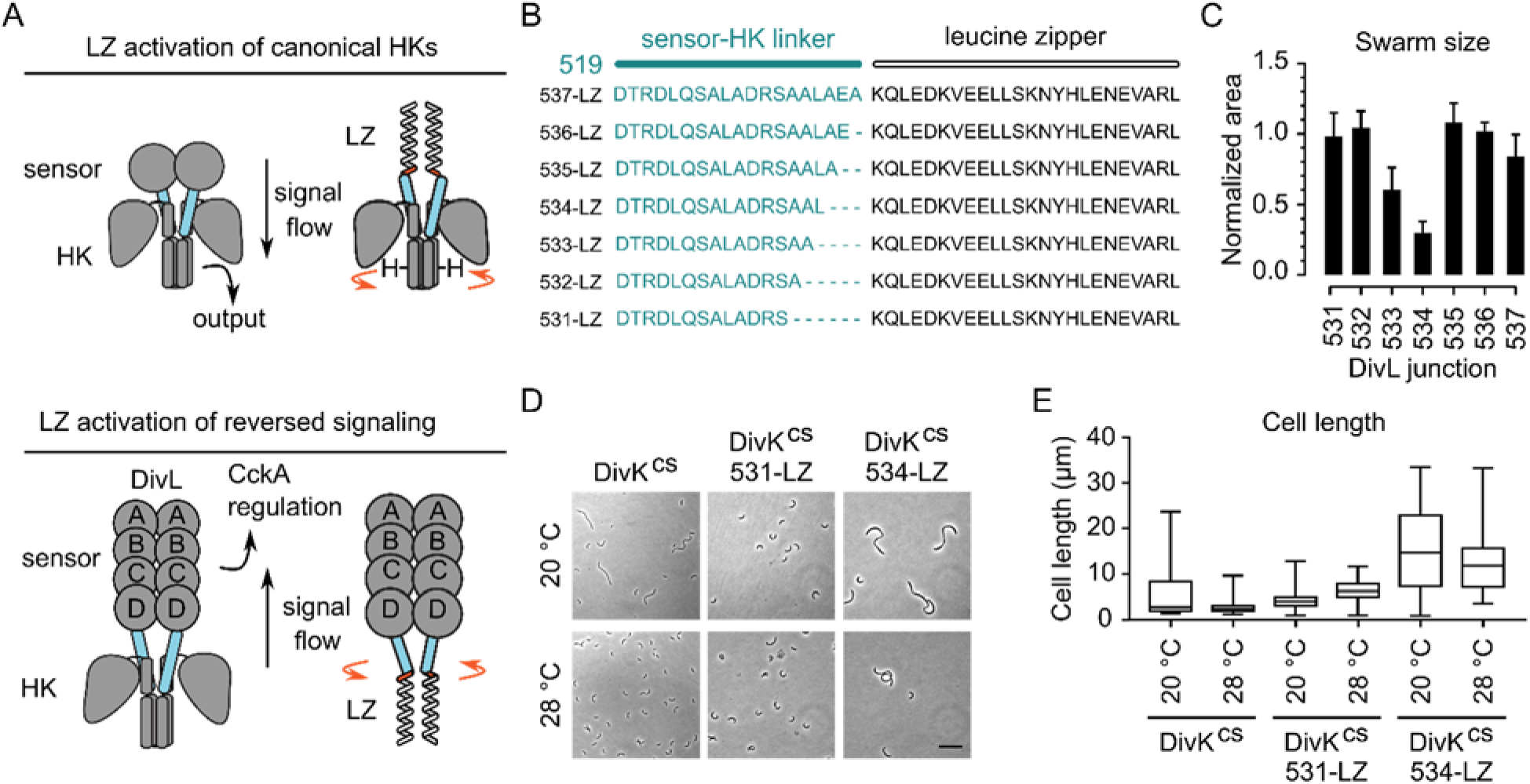
The leucine zipper fusion method was utilized to probe reverse signaling conformations and their effect on the cell cycle. (A) A schematic of the leucine zipper fusion method as employed in two previous studies of canonical histidine kinases and how it is applied in this study to modulate DivL in reverse. The fusion of the leucine zipper to the native coiled-coil of the dimerization/histidine phosphorylation domain determines the position of the histidine in the coiled-coil, resulting in periodic accessibility for phosphotransfer. (B) The sequences of the DivL-leucine zipper fusions used in this study. The coiled-coil linker of DivL was truncated from residue 537 while the leucine zipper sequence was maintained. (C) Quantification of the motility assay of *C. crescentus* strains expressing DivL-M2 leucine zipper fusions. *C. crescentus* were stabbed into 0.3% PYE agar supplemented with 0.3% xylose and incubated at 28 ºC for 3 days. Error bars represent SD (n=3). (D) Phase microscopy to visualize the cell phenotypes of DivL leucine zipper fusions expressed in the DivK^CS^ strain. Strains were grown overnight in PYE medium and diluted in PYE supplemented with 0.03% xylose to induce overexpression. Cells were grown again overnight at 28 or 20 °C. Scale bar denotes 10 µm. (E) Quantification of cell lengths for each condition. The center line is the median and the box extends to the 25th and 75th percentiles. The whiskers lie at the minimum and maximum values. N>18 cells for all samples.

### The leucine zipper periodically modulates cell cycle activity

Upon overexpression of each of the DivL-LZ fusions in *C. crescentus*, confirmed by Western blot (Figure S1), we observed that the swarm area changed periodically from 30-108% of wild-type as a function of the DivL linker length (Figure 4C, S6). This hallmark pattern indicates that a specific coiled-coil interface mediates swarm size reduction, while other coiled-coil interfaces have dampened or no impact on swarm size. Remarkably, the synthetic leucine zipper activation strategy indicates that the sensory helix can signal in reverse to modulate DivL’s *in vivo* regulatory functions and does not specifically require DivL’s HK domain.

Due to the distinct activity of each of the DivL-LZ fusions in the swarm assay, we hypothesized that select fusions may mimic the DivL-DivK~P binding conformation or mimic a CckA kinase activating conformation. If this is the case, we predicted that some of the DivL-LZ fusions would be able to recover the phenotype that results when DivL is unable to bind DivK~P. We used a DivK cold-sensitive (DivK^cs^) genetic strain (41, 46, 47) in which DivK D90G exhibits decreased binding to DivL (12). Growth of the DivK^cs^ strain at 20 °C resulted in significantly longer cells, with 15% of cells longer than 10 µm, than at the permissive temperature 28 °C, with no cells longer than 10 µm (Figure 4D-E), as previously reported (46, 47). Supplementing this strain with DivL-531-LZ resulted in less than 1% of cells with lengths longer than 10 µm at 20 °C (Figure 4D-E). In contrast, supplementing the DivK^cs^ strain with DivL-534-LZ caused severe cell division phenotypes (greater than 50% of cells longer than 10 µm) at both temperatures (Figure 4D-E). The DivL-531-LZ variant was able to rescue the DivKcs phenotype under cold temperature conditions, but the DivL-534-LZ variant was not. These results suggest that DivL-531-LZ may mimic the DivK~P binding conformation, but DivL-534-LZ may promote CckA phosphorylation in a manner similar to the DivKcs strain. Interestingly, we observed that DivL-531-LZ was functional in the DivKcs background, while yielding no impact in the wild-type background. This suggests that dynamic feedback regulation mediated through DivK may be able accommodate the functional impact of DivL-531-LZ.

### The leucine zipper periodically modulates localization of DivL

Since we observed that the HK domain impacts DivL’s subcellular pattern (Figure 2C-D), we analyzed the localization pattern of the DivL-LZ chimeric library in *C. crescentus* (Figure 5A-B). Strikingly, a periodic trend of cell pole accumulation emerged as a function of the DivL linker length (Figure 5A-B). This result suggest that conformational changes mediated by the sensor-HK linker could regulate DivL’s subcellular localization pattern. In designing the leucine zipper approach, we hypothesized that the strong effect of the leucine zipper would require an intact PAS signal transmission motif. We thus introduced D516A and T518V mutation into the DivL-532-LZ chimera, which had a largely diffuse population (Figure 5A-C). In comparison, the signal transmission mutants of DivL-532-LZ re-gained the ability to form foci (Figure 5D-E). This result indicates that the high affinity leucine zipper dimerization can only exert an effect upon DivL subcellular localization with an intact PAS signal transmission motif. As a whole, the leucine zipper DivL localization data also support our hypothesis that DivL localization can be regulated by reversed signaling originating from the pseudo-HK domain.

**Figure 5:**
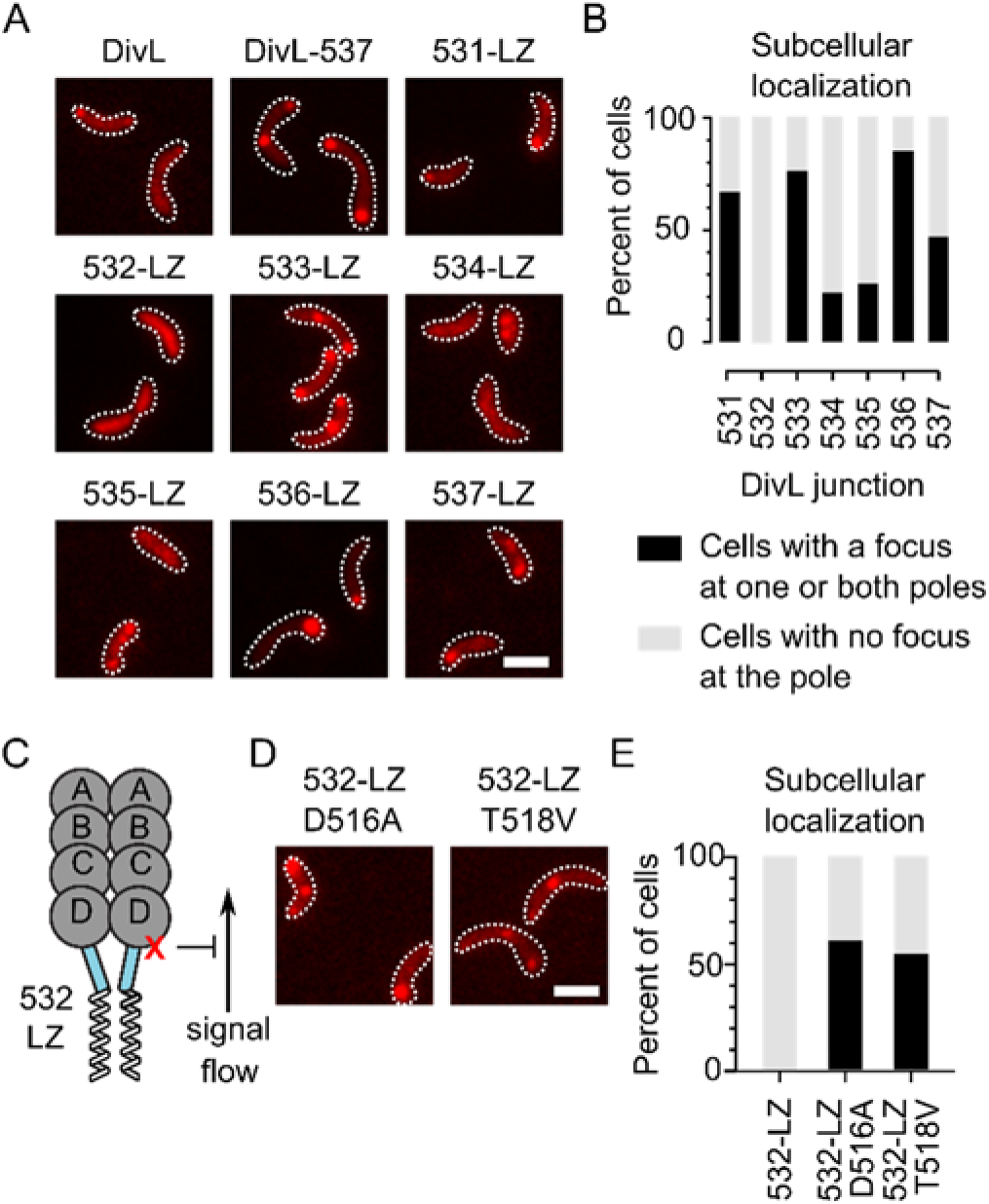
The leucine zipper synthetically and periodically modulates the localization of DivL and requires an intact signal transmission motif. (A) Fluorescence microscopy to visualize the subcellular localization of DivL-mCherry leucine zipper fusions expressed in *C. crescentus*. The variants were induced from the chromosomal xylose promoter in M2G supplemented with 0.03% xylose for 4 hours. Scale bar denotes 2 µm. (B) Quantification of the percent of cells with the DivL-LZ variants localized at the cell pole. N>97 cells for all samples. (C) A model of DivL’s sensor domain fused to the leucine zipper with the signal transmission motif mutations indicated by an X. (D) Fluorescence microscopy to visualize the subcellular localization of DivL-mCherry leucine zipper fusion mutants expressed in *C. crescentus*. The variants were induced from the chromosomal xylose promoter in M2G supplemented with 0.03% xylose for 4 hours. (E) Quantification of the percent of cells with the DivL-LZ variants localized at the cell pole. N>54 cells for all samples.

A possible explanation for both the DivL-LZ fusions that were inactive in the swarm assay and the ones that did not form foci is that they form misfolded aggregates. We chose two DivL-LZ fusions to purify and analyze by gel filtration (Figure S7), each having a predicted monomeric weight of 58 kDa. Both DivL-532-LZ, which was inactive in the swarm assay and had diffuse localization, and DivL-536-LZ, which was also inactive in the swarm assay but exhibited the most polar localization eluted as a single peak with an apparent molecular weight of approximately 86 kDa. The results indicate that the DivL-LZ fusions have identical oligomerization profiles as well-folded proteins.

### DivL’s PAS-B and PAS-D are required for DivL mediated regulation of cell motility

The leucine zipper experiments have revealed that DivL transmits signal through a conformational change into its repurposed sensory effector domain. We have also identified two outputs of the effector domain: regulation of swarm size and subcellular localization. The N-terminal sensor domain comprises a transmembrane region followed by a domain of unknown function, DUF3455, referred to hereafter as Domain A. Homology structure prediction did not result in any statistically significant hits for this domain, and the secondary structure prediction is distinct from the PAS fold and other known sensory domains (Figure S8A-B). The remaining three sensory domains display have homology with the PAS domain, and these have been referred to as PAS B, PAS C, and PAS D. We individually deleted each of these domains, confirmed expression by Western blot analysis (Figure S1), and evaluated the effect of DivL variant overexpression upon swarm size (Figure 6A, S9). Deletion of PAS B resulted in a smaller swarm size with an average area of 24% ± 3% of the empty vector control. In contrast, deletion of PAS D resulted in a larger swarm size with an average area of 90% ± 1% of the empty vector control. Overexpression of DivL-ΔA (Δ28-133) and DivL-ΔPAS C did not result in significantly different swarm areas compared to wild-type DivL. These results indicate that PAS B and PAS D are involved in regulating swarm motility. Identification of PAS B’s involvement in cell cycle regulation is consistent with a previous biochemical study that showed that DivL and DivL-ΔA can inhibit CckA autophosphorylation, but DivL-ΔAB cannot (25).

**Figure 6:**
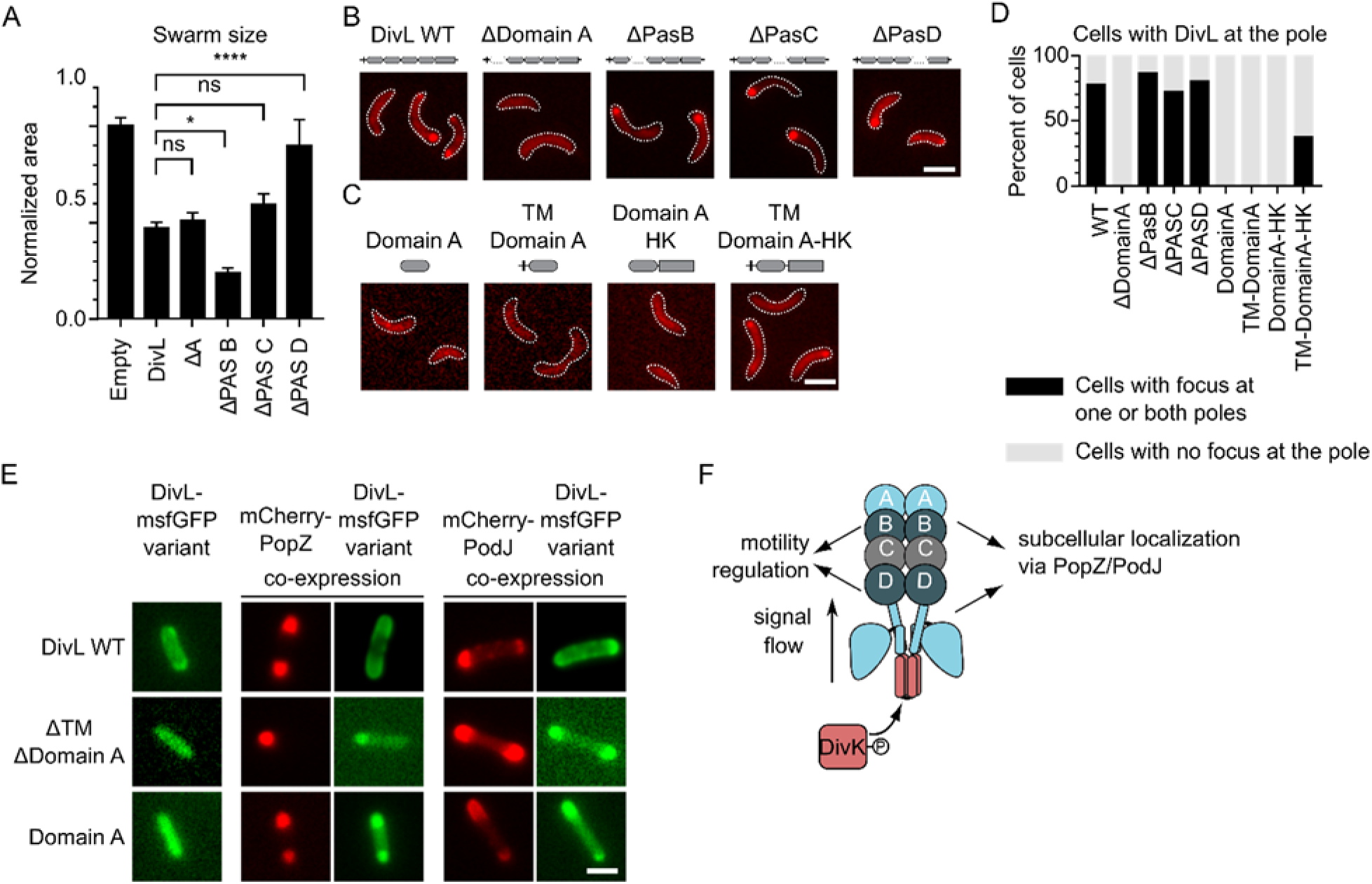
DivL’s PAS-B and PAS-D are required for DivL mediated regulation of cell motility and Domain A and the HK of DivL regulate cell-pole binding through scaffold interactions. (A) Quantification of the motility assay of *C. crescentus* strains expressing DivL-M2 domain deletions. *C. crescentus* were stabbed into 0.3% PYE agar supplemented with 0.3% xylose and incubated at 28 ºC for 3 days. Error bars represent SD (n=3). Significance was determined with a Dunnett’s multiple comparisons test by comparing the mean area of each strain to the mean area of DivL-WT (ns: P>0.05, *: P ≤ 0.05, **: P ≤ 0.01, ***: P ≤ 0.001). (B) Fluorescence microscopy to visualize the subcellular localization of DivL-mCherry domain deletion variants and (C) a panel of potential minimal localization constructs in *C. crescentus*. The variants were induced from the chromosomal xylose promoter in M2G supplemented with 0.03% xylose for 4 hours. Scale bar denotes 2 µm. (D) Quantification of the percent of cells with DivL localization at the cell pole. N>100 cells for all samples. (E) Heterologous expression of DivL-msfGFP variants with mCherry-PopZ or mCherry-PodJ in *E. coli*. DivL variants were expressed from the pBAD plasmid using 10 mM arabinose for 2 hours. mCherry-PopZ and mCherry-PodJ were expressed from the pACYC plasmid using 0.05 mM IPTG for 2 hours. Scale bar denotes 2 µm. (F) Model of DivL’s signaling flow with the DivK binding site shaded red and the domains that impact swarm motility shaded dark blue and localization shaded light blue.

### Domain A and the HK of DivL regulate cell-pole binding through scaffold interactions

We next analyzed the role of each DivL domain towards DivL subcellular localization. Each domain deletion variant could form foci, with the exception of DivL-ΔA (Δ28-133) that was unable to accumulate as focus (Figure 6B, D). We asked if Domain A was sufficient for foci formation or polar localization, but found that when this domain was expressed with or without the transmembrane domain, it remained diffuse. Considering the role that the HK seemed to play in localization at the poles (Figure 2C-D) (8, 12), we fused Domain A directly to the HK and found that localization at the poles was restored (Figure 6C-D). The transmembrane domain was required for this minimal construct to localize to the cell pole (Figure 6C-D). We also observed that a second variant that lacks a helical portion that may be an input coil into PAS B, DivL(Δ28-170) (Figure S10A, C) was unable to form foci within the cell. Collectively, these results suggest that cell pole accumulation minimally requires Domain A, the transmembrane domain, and the HK domain (Figure 6B-D S10A, C). To further probe the role of dimerization Domain A(1-170) in localization, we fused a leucine zipper to residue 170 and observed irregular protein aggregation that was located in the cell body in 92% of cells (Figure S10B-C). These results suggest that dimerization of Domain A alone is insufficient to promote DivL foci formation. Recalling that DivL-518, which could also be called DivL-ABCD, displayed enhanced localization at the poles (Figure 2C-D), we constructed several PAS domain combinations and found that no combination was able to localize similar to DivL-WT (Figure S10B-C). From this domain analysis, we conclude that interactions between Domain A and the HK may regulate DivL’s localizations pattern.

We next investigated whether Domain A mediates DivL’s cell pole localization through protein-protein interactions with two key cell-pole scaffolds: PodJ or PopZ. PodJ has been shown to recruit DivL to one cell pole in *C. crescentus* (48) while PopZ and DivL have been shown to interact in a heterologous *E. coli* reconstitution assay (49). We performed a similar assay in which we expressed DivL variants with either PodJ or PopZ in *E. coli* and measured their co-localization as an indication of interaction (Figure 6E, S10D). Wild-type DivL co-localized with PopZ and to a lesser extent PodJ. We observed that DivL Domain A alone co-localized with both PopZ and PodJ. Expression of wild-type DivL and Domain A disrupted the typical monopolar localization pattern of PopZ in *E. coli* indicative of an interaction between DivL and PopZ (49). We also observed that DivL-ΔTMΔA was able to co-localize with PopZ and PodJ. This result was unexpected based on the diffuse localization of DivL-ΔA in *C. crescentus* (Figure 6B, D), however is consistent with earlier studies that have suggested a cell-pole binding site at the C-terminus of DivL (21, 43, 44). The *E. coli* reconstitution assay results demonstrate that DivL mediates interaction with PopZ and PodJ through both domain A and likely the C-terminal residues of the HK domain (Figure 6F).

## Discussion

Kinase-kinase signaling cross-talk may be wide-spread throughout bacteria, and our work examining DivL-CckA has revealed that cross-talk can be mediated upon reversed signaling. In this model stimulation of the HK domain results in a conformational change in the sensor helix (Figure 2) and through the signal transmission motif (Figure 3), ultimately modulating CckA activity via PAS-B and PAS-D (Figure 6A), and changes in DivL subcellular accumulation via Domain A (Figure 6B-E). A core strategy for directly testing the reverse signaling model, was use of a leucine zipper fusion to synthetically stimulate reverse signaling (Figure 4–5). Using this strategy, we successfully demonstrated that DivL-531-LZ can mimic the DivL-DivK~P bound state to repress CtrA phosphorylation in the DivK^cs^ background that encodes a DivK variant that binds DivL poorly (Figure 4D-E). We anticipate that further biochemical studies of the DivL-LZ fusions in CckA kinase activity assays will help us to further characterize DivL’s allosteric activator role.

Here we have discovered direct interactions between two scaffolds and DivL’s Domain A. This raises several questions remain about the inter-relationship between Domain A and the HK. For example, does the HK regulate Domain A’s ability to bind the scaffolds through a conformational change induced by DivK~P binding? Is there a direct interaction between Domain A and the HK that creates a new scaffold binding surface? Moreover, it also raises a set of questions about the impact of scaffold binding to DivL upon CckA regulation. For example, do PopZ and PodJ’s scaffold interactions with Domain A regulate the CckA kinase activity in a "forward" signaling manner? Or do PopZ and PodJ merely serve as benign scaffolds that regulate DivL-CckA subcellular position?

To interrogate this reversed signaling flow we applied mutations that "short circuit" signal transmission from PAS signaling domains together with a leucine zipper approach to synthetically stimulate reverse signaling. Using DivL and the asymmetric division multi-kinase network as a case-study, here we demonstrated that this leucine zipper synthetic activation strategy (Figure 4A-C) may provide a broadly useful tool to map-out signaling flux in bacterial multi-kinase networks. Moreover, "short circuit" mutations to the highly conserved PAS signal transmission motifs (Figure3) provide a powerful companion approach to interrogate the function of PAS sensors. Beyond histidine kinases, coiled-coil linkers frequently connect signaling domains within bacterial signaling proteins including ATPases, phosphatases, cyclic nucleotide cyclases, HAMP signaling domains, and diguanylate cyclases (50).

Studies of pseudoenzymes have highlighted functions that extend beyond canonical enzymatic activity (22, 51, 52). The functions of these pseudokinases can be described in four categories, three of which DivL is proposed fulfill in *C. crescentus*: modulator (8, 12), competitor, anchor (8), and signal integrator (5, 12). More broadly, studies of the DivL pseudokinase highlight the possibility of a new category of pseudoenzyme regulation that include a ligand binding domain that regulates an enzymatic domain in a "forward signaling" manner. For example, a similar reverse signaling as we have observed with DivL, has also been observed with the *C. crescentus* signaling protein PopA (53). For canonical members of the diguanylate cyclase protein family, phosphorylation of an N-terminal receiver domain regulates a C-terminal catalytic domain’s cyclization of GTP into the secondary messenger cyclic-di-GMP (c-di-GMP). In contrast, PopA utilizes a catalytically dead C-terminal domain that has been repurposed as a c-di-GMP sensory domain. C-di-GMP binding to the pseudo-catalytic domain enables the receiver domain to interact with downstream targets to regulate the cell cycle (53). Thus, DivL and PopA provide two examples of pseudoenzymes that re-wire the input-output assignments and use reverse signaling flow. In the case of DivL this includes a functional chimera of a pseudokinase domain that regulates a "pseudosensory" domain as a repurposed effector domain. We speculate that the DivL reversed signaling model may indicate broader simultaneous repurposing of catalytic domains as sensors and sensory domains as output effector domains amongst pseudoenzymes.

## Methods

### Construction of Plasmids and Strains

All experiments were performed using *Caulobacter crescentus* NA1000 (also known as CB15N) and *Escherichia coli* DH5α (Invitrogen) and BL21 (Novagen). *C. crescentus* NA1000, WSC11311 *DivK341* (DivK^cs^), and the φCR30 phage were kind gifts from Dr. Lucy Shapiro (Stanford University School of Medicine). *E. coli* BL21 strains containing the pBAD-DivL-msfGFP and the pACYC-mCherry-PopZ plasmids were generously given by Dr. Grant Bowman (University of Wyoming Department of Molecular Biology). DNA oligos, plasmid construction methods, plasmids, and strains used in this study are further described in the Supplemental Text and listed in Table S1-S5.

### Growth Conditions and Inducer Concentrations

*C. crescentus* strains were grown at 28°C in PYE (peptone yeast extract) or M2G (minimal medium supplemented with glucose) (54). *E. coli* strains used for protein purifications were grown at experiments were grown at 28 °C in LB medium and for microscopy experiments were grown at 37 °C in LB medium unless otherwise stated. When required, protein expression was induced by adding 0.002-0.5 mM Isopropyl β-D-1-thiogalactopyranoside (IPTG) or 0.5-10 mM arabinose in *E. coli*, and 0.003%–0.3% xylose or 0.05-0.5 mM vanillic acid in *C. crescentus*. The induction time for microscopy experiments is usually 2 hours in *E. coli* and 4 hours in *C. crescentus*.

### Swarm Plate Assay

Cells were grown to mid-log phase overnight in PYE medium and the appropriate antibiotic. Cells were normalized by dilution in PYE medium to the culture with the lowest OD600. Cells were stabbed into 0.3% PYE agar with the appropriate antibiotic and 0.3% xylose in 15 cm diameter culture plates using a Boekel replicator. Plates were incubated at 28 °C for 3 days. Plates were visualized using a ChemiDoc XRS+ system (Bio-Rad). Three replicate plates were analyzed. Swarm area was measured using ImageJ (55). Swarm areas were normalized to the empty vector control on each plate. The error for the empty vector control was calculated by dividing the standard deviation of the areas by the average area. The error for the experimental areas was calculated as the standard deviation of the areas after normalization to the empty vector control. Significance was determined using Prism (GraphPad) with a Dunnett’s multiple comparisons test by comparing the mean area of each strain to the mean area of DivL-WT (ns: P>0.05, *: P ≤ 0.05, **: P ≤ 0.01, ***: P ≤ 0.001).

### Fluorescence Microscopy

*C. crescentus* cells were grown in M2G medium with the appropriate antibiotics at 28 °C overnight to OD600=0.2-0.8. DivL-mCherry variants were induced with 0.03% xylose for 4 hours. Cells were diluted as needed an immobilized on a 1.5% agarose in M2G pad. *E. coli* cells were grown in LB medium with the appropriate antibiotics at 37 °C and diluted to OD600 = 0.2. DivL variants were expressed from the pBAD plasmid using 10 mM arabinose for 2 hours. mCherry-PopZ and mCherry-PodJ were expressed from the pACYC plasmid using 0.05 mM IPTG for 2 hours.

Phase microscopy was performed by using a Nikon Eclipse T*i*-E inverted microscope equipped with an Andor Ixon Ultra DU897 EMCCD camera and a Nikon CFI Plan-Apochromat 100X/1.45 Oil objective and intermediate 1.5x magnification. DIC (differential interference contrast) microscopy was performed with a Nikon CFI Plan-Apochromat 100X/1.45 Oil DIC objective with a Nikon DIC polarizer and slider in place.Carl Zeiss Immerson Immersion Oil 518 was used. Excitation source was a Lumencor SpectraX light engine. For single-channel experiments, mCherry was visualized with the DAPI/GFP/TRITC Chroma filter cube. For two-channel experiments, mCherry was visualized with the CFP/YFP/MCHRY MTD TI Chroma filter cube and the DAPI/GFP/TRITC Chroma filter cube was used to image EGFP (470/40X) and sfGFP (515/30M). Images were acquired with Nikon NIS-Elements AR software.

### Fluorescence Image Analysis

ImageJ (55) was used to adjust LUT, pseudocolor, and crop images. Image J Cell Counter and Nikon-NIS Elements AR were used to manually count cells and foci. ImageJ and MicrobeJ (56) were used for manual cell counting, cell length analysis, and two-channel correlation calculations.

### Multiple Sequence Alignments

The DivL homolog sequences (57) were aligned using ClustalΩ in the MPI Bioinformatics Toolkit (58). The phylogenetic tree was calculated in Jalview using BLOSUM62 Average Distance (59). Protein visualization was performed with USCF Chimera (60).

### Structure Predictions

Transmembrane regions were predicted using TMpred (61). Domain A homology with known protein folds was calculated using HHpred, and the DivL PAS D signal transmission motif was modelled to YF1 using MODELLER (62). PSIPRED (63) was used to predict the protein secondary structure of Domain A.

## Acknowledgements

We thank Dr. Lucy Shapiro for providing critical *C. crescentus* strains and Dr. Grant Bowman for providing critical *E. coli* strains that supported this study.

