## Supplemental Information for "Reversed Signaling Flow of a Bacterial Pseudokinase"

*W. Seth Childers

**This PDF file includes:**

Figures S1 to S10
Supplementary Materials and Methods
Tables S1 to S5
SI References

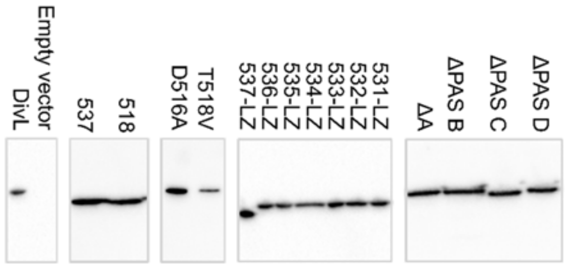

Figure S1: **Western blot analysis of overexpression of DivL variants** Western blots were performed to confirm overexpression of DivL variants in the strains used in the swarm assays. The different blots were scaled relative to the protein ladder.

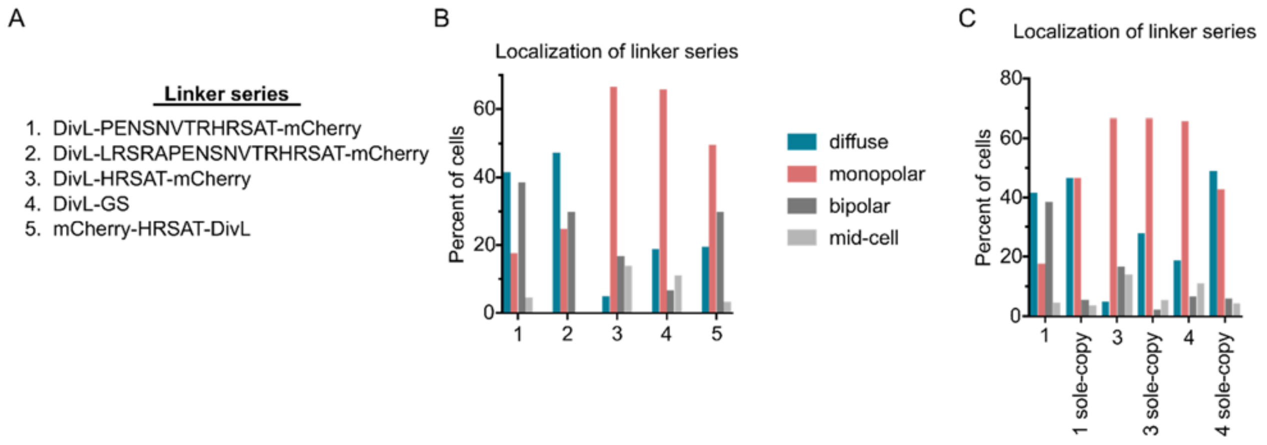

Figure S2: **The DivL fluorescent protein affects subcellular localization.** (A) The designs and sequences of the mCherry and DivL linker series. (B) Fluorescence microscopy was used to visualize the subcellular localization of DivL-mCherry fusions expressed in *C. crescentus*. The variants were induced from the chromosomal xylose promoter in M2G supplemented with 0.03% xylose for 4 hours. The localization pattern differs depending on the design and linker sequences. (C) The variants were expressed in a *divL* deletion strain and the localization patterns were analyzed. Upon deletion of wild-type DivL, more cells exhibited diffuse localization in each case, but the localization still varied based upon the design and linker sequences of the fusions. Monopolar foci were almost exclusively found at the stalked pole in all strains. N>115 cells for all samples.

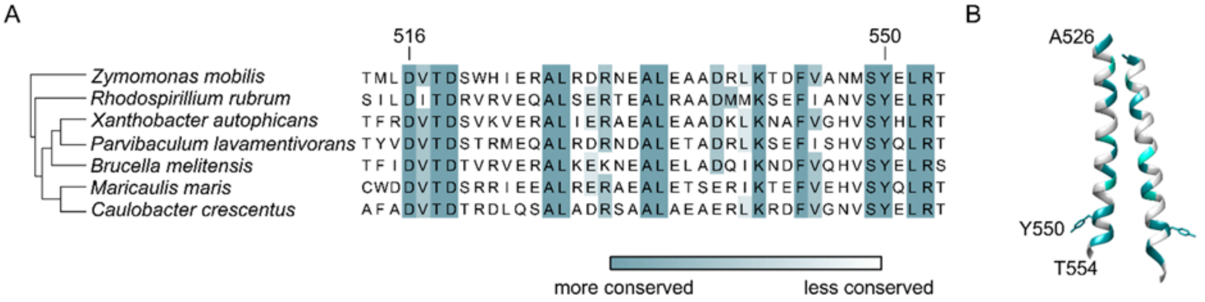

Figure S3: **The DivL sensory linker is conserved at the coiled-coil interface.** (A) Alignment of the sensory helix of DivL homologs in the alpha-proteobacteria. More highly conserved residues are depicted by darker shades. The alignment maintains the tyrosine in the phosphorylatable position (Y550 in C. crescentus) and shows that the signal transmission motif DVT or DIT and coiled-coil interfaces are conserved across the class (D516-T518 in *C. crescentus*). (B) The structure of the sensory helix-DHp of C. crescentus DivL (PDB: 4Q20) (1) from residues A526-T554. The structure of the tyrosine in the phosphorylatable position Y550 is shown. Conservation is depicted in colors according to the alignment in Figure 2E.

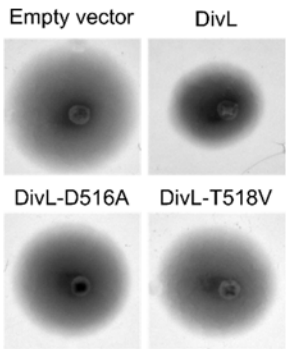

Figure S4: **The conserved signal transmission motif is critical for DivL swarm activity.** Motility assay of C. crescentus strains expressing DivL-M2 point mutations to the signal transmission motif. Swarm areas are quantified in Figure 3C.

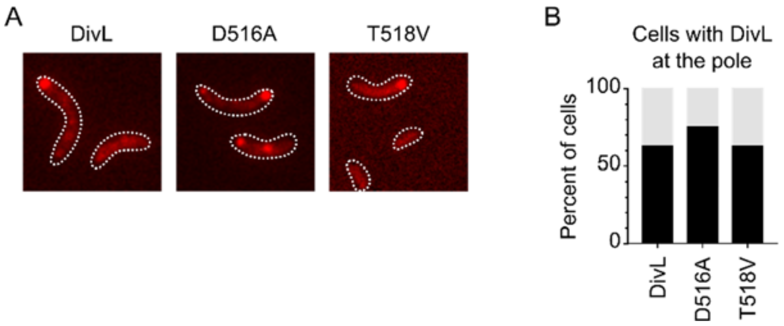

Figure S5: **A DivL signal transmission motif mutations have a mild effect on subcellular localization.** (A) Fluorescence microscopy to visualize the subcellular localization of the DivL-mCherry signal transmission motif mutants expressed in *C. crescentus*. The variants were induced from the chromosomal xylose promoter in M2G supplemented with 0.03% xylose for 4 hours. (B) Quantification of the percent of cells with the DivL variants localized at the cell pole. N>110 for all samples.

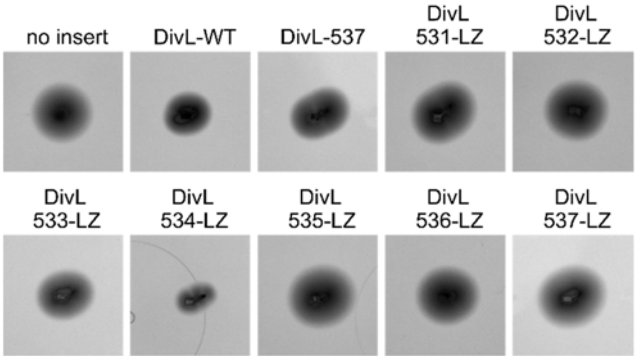

Figure S6: **The leucine zipper fusion synthetically modulates the effect of DivL on the cell cycle** Motility assay of C. crescentus strains expressing DivL-M2 leucine zipper fusions. Swarm areas are quantified in Figure 4D.

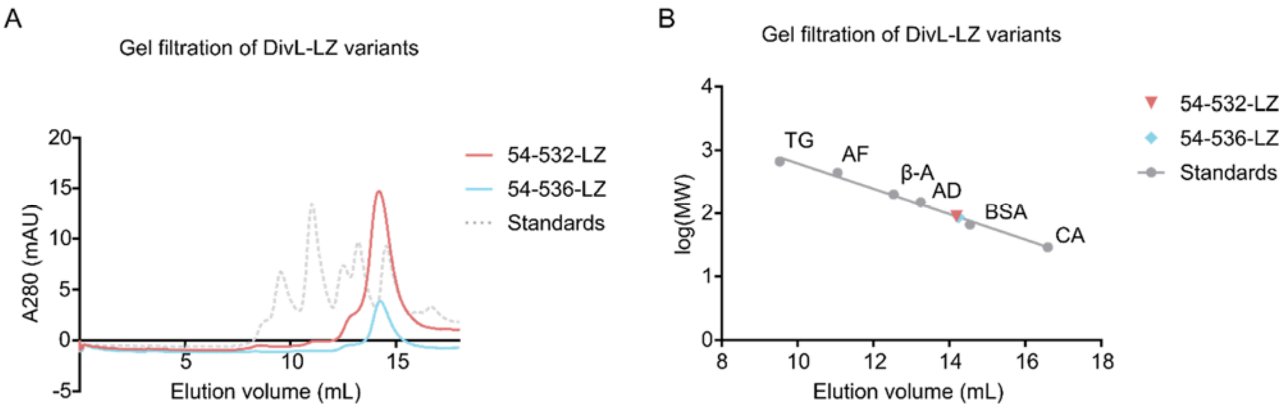

Figure S7: **Gel filtration analysis of two DivL-LZ variants** (A) Gel filtration elution curve of the protein standards (gray) and DivL-54-532-LZ (red) and DivL-54-536-LZ (blue) shows that both leucine zipper fusions were eluted with the same volume of buffer. (B) A standard curve was created to calculate the apparent molecular mass of the leucine zipper fusions. The protein standard sizes were as follows: thyroglobulin (TG, 669 kDa), apoferritin (AF, 443 kDa), β-amylase (β-A, 200 kDa), alcohol dehydrogenase (AD, 150 kDa), bovine serum albumin (BSA, 66 kDa), and carbonic anhydrase (CA, 29 kDa).

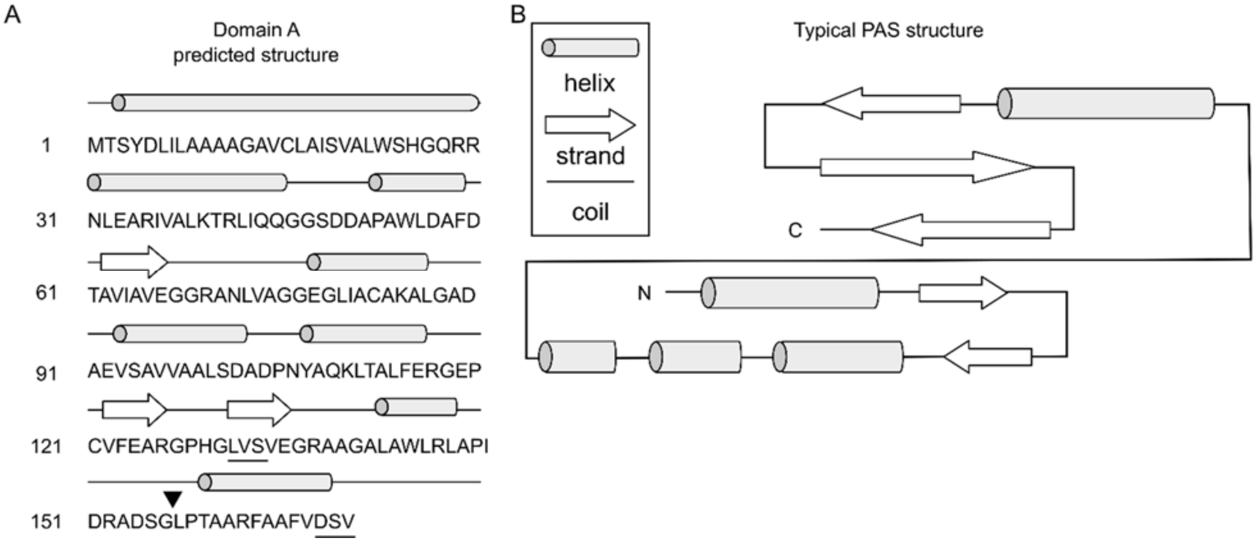

Figure S8: **The DivL DUF Domain A does not have a predicted PAS-like structure.** (A) The secondary structure of Domain A was predicted based upon primary sequence by PSIPRED (2). The arrow points to the designation of the end of DUF3455. The underlined residues indicate the signal transmission motif-like sequences that dictated the 133 and 170 cut-offs. (B) Representative topology of the PAS domain, adapted from (3).

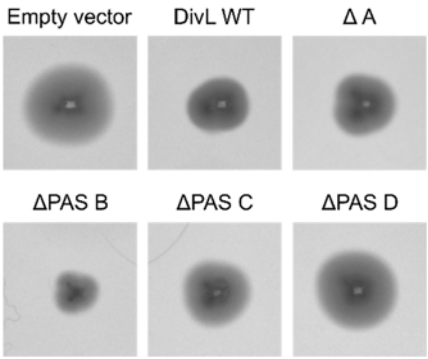

Figure S9: **DivL's PAS-B and PAS-D are required for DivL mediated regulation of cell motility.** Motility assay of C. crescentus strains expressing DivL-M2 domain deletions. Swarm areas are quantified in Figure 6A.

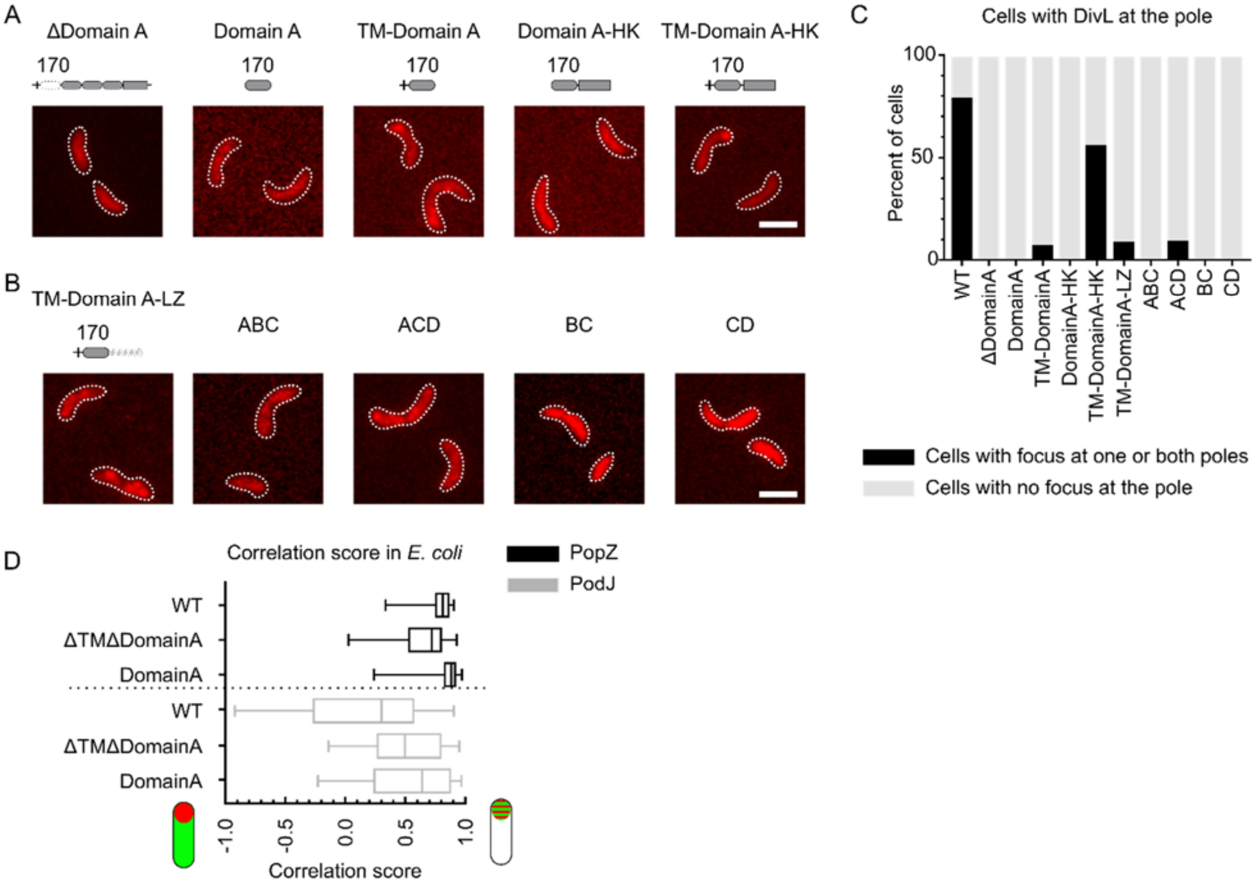

Figure S10: **DivL's Domain A and HK regulate cell-pole binding.** (A) Fluorescence microscopy to visualize the subcellular localization of the DivL-mCherry Domain A(170) constructs expressed in *C. crescentus*. (B) Fluorescence microscopy to visualize the subcellular localization of the DivL-mCherry Domain A(1-170)-LZ and PAS domain combinations. The variants in (A) and (B) were induced from the chromosomal xylose promoter in M2G supplemented with 0.03% xylose for 4 hours. (C) Quantification of the percent of cells with the DivL variants in (A) and (B) localized at the cell pole. N>36 cells for all samples. (D) Quantification of co-localization between the DivL variants and PopZ or PodJ in Figure 7C. A correlation score of +1 indicates complete co-localization and a score of -1 indicates no co-localization. The center line is the median and the box extends to the 25th and 75th percentiles. The whiskers lie at the minimum and maximum values. N>72 cells for all samples.

**Supplementary Information Text**

**SI Materials and Methods**

**Plasmid construction.** Restriction enzymes were purchased from Thermo Scientific or Invitrogen. PCR reactions were performed in 50 µL reaction mixtures containing 3% (v/v) DMSO, 1.3 M betaine, 0.3 µM each primer, and 0.2 mM each dNTP, and 1U Phusion High-Fidelity DNA Polymerase (Thermo Scientific). Gibson assembly (4) reactions were performed in 20 µL with 100 ng backbone and typically a 1:5 backbone:insert ratio, with 0.08U T5 Exonuclease (New England Biolabs), 1U Phusion High-Fidelity DNA Polymerase (Thermo Scientific), and 80U Taq DNA Ligase (New England Biolabs). An annealing temperature of 55 °C was used for most reactions. Plasmids and primers were designed using the J5 device editor software (5). Oligonucleotides were synthesized by IDT (Coralville, IA) and all DNA sequencing reactions were performed by The University of Pittsburgh Genomics Research Core or Genewiz (South Plainfield, NJ). DNA oligos, plasmid construction methods, plasmids, and strains used in this study are listed in Table S1-S5.

**DivL-mCherry variants.** Plasmid WSC10153-pXCHYC6-divL was made by Gibson assembly of full-length divL into pXCHYC-6 (6) and served as the template for all further DivL plasmid designs. Full length divL was amplified from the template cosmid 2G1 (7) using primers KAK1 and KAK2. pXCHYC-6 was digested with SacI. This backbone and the insert were assembled using the Gibson assembly method (4) resulting in an integrating plasmid at the C. crescentus chromosomal xylose locus encoding C-terminal mCherry-tagged DivL. C. crescentus cells were transformed by electroporation. Briefly, cells were grown in PYE medium overnight and rinsed three times with cold sterile water. Cells were resuspended in water and 80 µL of cells plus 10 µL of plasmid were electroporated with a time constant of 3.5-5.0 ms. Colonies were screened for integration at the xylose locus using RecUni-1, a primer that anneals to the plasmid, and RecXyl-2, a primer that anneals to the 5’ region of the chromosomal promoter (6).

DivL-LZ fusion variants: The leucine zipper sequence was embedded into the reverse primer with overhangs overlapping the DivL truncation and the backbone. PCR was used to amplify the DivL-LZ fusion, and Gibson assembly was used to insert the fusion into the PXCHYC-6-SacI or PCR-amplified pBXSPA-2 backbones described in these methods.

**DivL-mCherry linker panel.** Cloning plasmid pWSC10153 resulted in the linker PENSNVTRHRSAT between DivL and mCherry. To vary this linker sequence, primers were used to amplify the original pXCHYC-6 backbone so that the DivL sequence would align with different regions of the multiple cloning site. The primers used to amplify the backbones and DivL inserts for this series are found in Table S1.

**Table S1. Plasmid design of DivL-mCherry linker panel**

| Plasmid | Description | Forward primer backbone | Reverse primer backbone | Forward primer DivL | Reverse primer DivL |
|---|---|---|---|---|---|
| pKAK0141 | pXCHYC-6 DivL LRSRAPENSNVTRHRSAT mCherry linker | KAK173 | KAK174 | KAK175 | KAK176 |
| pKAK0142 | pXCHYC-6 DivL HRSAT mCherry linker | KAK177 | KAK178 | KAK179 | KAK180 |
| pKAK0143 | pXCHYC-6 DivL GS mCherry linker | KAK181 | KAK182 | KAK183 | KAK184 |
| pKAK0145 | pXCHYN-6 DivL mCherry HRSAT linker | KAK189 | KAK190 | KAK191 | KAK192 |

**Phage transduction.** Single-copy DivL-mCherry strains were made by using ϕCR30 phage transduction (8) to move the streptomycin divL gene replacement from WSC0427 (9) to the strains containing the divL-mcherry copy under the xylose locus. To prepare ΔdivL phage, WSC0427 was grown in 2 mL of PYE medium supplemented with streptomycin and spectinomycin to stationary phase. Three dilutions of ϕCR30 phage in 5 µL was added to 0.5 mL of culture and incubated at room temperature for 15 minutes. The mixture was plated in top agar on a plain PYE plate and incubated at 28 °C overnight. The phage plaques were resuspended in 5 mL PYE and 100 µL chloroform, incubated for 10 minutes at room temperature, followed by centrifugation for 30 minutes at room temperature at 8,000 RPM.

As an example, to transduce the KAK43 strain, it was grown in 2 mL PYE supplemented with chloramphenicol overnight. Expression of DivL-mCherry was induced with 0.3% xylose for 2 hours. 100 µL of the phage prepared above was mixed with 0.5 mL cells and incubated at room temperature for 40 minutes. Cells were centrifuged at 15,000 g for 2 minutes and resuspended in 0.5 mL of PYE medium, then incubated with shaking at 28 °C for 2 hours. Cells were plated on PYE agar supplemented with 0.3% xylose, spectinomycin, and streptomycin. Colonies were screened for insertion of the streptomycin replacement at the divL locus using the primers KAKqc0046 that binds downstream of the divL locus and KAKqc0049 that binds in the streptomycin cassette. This resulted in strain KAK115: ΔdivL with pWSC10153 DivL-mCherry integrated at the xylose locus.

**DivL point mutants.** DivL point mutants were designed as two-fragment Gibson assemblies with the point mutant embedded in the fragment amplification primers. pKAK0009-pXCHYC6-SacI-DivL-D516A was made by amplifying Insert 1 using primers KAK1 and KAKqc0019 and Insert 2 using primers KAKqc0018 and KAK2 from plasmid WSC10153 as a template. pXCHYC-6 was digested with SacI. This backbone and the inserts were assembled using the Gibson assembly method resulting in an integrating plasmid at the C. crescentus chromosomal xylose locus encoding the C-terminal mCherry-tagged DivL mutant.

**DivL-M2 variants.** pBXMCS-2 backbone was amplified by PCR using the primers KAK9 and KAK10. Fragments were amplified from WSC10153 or an already existing construct in pXCHYC-6. The backbone and insert were assembled using the Gibson assembly method resulting in a C. crescentus high-copy plasmid for the xylose-inducible expression of the C-terminal M2-tagged DivL variant.

**pBAD-DivL-msfGFP variants.** The pBAD-DivL-msfGFP plasmid was isolated from a strain generously given by Dr. Grant Bowman (University of Wyoming Department of Molecular Biology) (10). The pBAD backbone, excluding DivL, was amplified using the primers KAK321 and KAK322. A divL fragment encoding 134-769 was amplified from plasmid WSC10153. The backbone and insert were assembled using the Gibson assembly method resulting in an E. coli arabinose-inducible plasmid encoding C-terminal msfGFP-tagged DivL 134-769.

**Western Blot.** Western blot analysis was used to determine protein levels of each DivL-M2 variant. Cells were grown in 2 mL PYE medium to early log phase at 28°C. Overnight induction with 0.3% xylose began at inoculation. 1 mL cells were harvested by centrifugation and resuspended in SDS 100 µL sample loading dye. Samples were treated by heating (75°C for 10 min). Samples volumes were normalized to the lowest OD600 upon collection and separated by 10% SDS–PAGE. Proteins were transferred onto a PVDF membrane (GE Healthcare). The membrane was blocked in 5% milk (AmericanBio) overnight at 4°C. The anti-M2 (FLAG) antibody (Sigma-Aldrich) (1:5000 in TBST for 1 hour at 4°C) (Bulldog Bio) was used with a goat anti-rabbit IgG-peroxidase secondary antibody (Sigma-Aldrich) (1:10,000 in TBST for 1 hour at room temperature). PVDF membranes were treated with an ECL western blotting kit (Thermo Scientific) and visualized using a ChemiDoc XRS+ system (Bio-Rad).

**Protein Purification:** Plasmids pKAK0137b and pKAK0137f were transformed into BL21 cells, and plated onto 100 µg/mL ampicillin LB plates and grown overnight at 37 °C. From a single colony, an overnight 10 mL 50 µg/mL ampicillin LB culture was inoculated and grown to saturation overnight. From this saturated culture two 1 L LB cultures were inoculated and grown to mid-log phase (0.6 OD). Expression of the DivL-LZ fusions was induced with 333 µM isopropyl-b-D-thiogalactopyranoside (IPTG) for 4 hours at 25 °C. The cells were harvested by centrifugation (4 °C for 20 minutes at 3,700 g). The resulting pellet was resuspended in 50 ml 50 mM HEPES pH=8, 0.5 M KCl and centrifuged (3,700 g at 4 °C for 20 minutes) to yield a cell pellet stored at -80 °C. Cells were thawed on ice and resuspended in 50 ml of lysis buffer (50 mM HEPES pH 8.0, 0.5 M KCl, 1 mM DTT, 25 mM imidazole, 10% glycerol, and 200 U of Benzonase Nuclease (Sigma) supplemented with SIGMAFAST protease inhibitor tablets (Sigma)). The cell suspension was lysed with three passes through the Emulsiflex at 20,000 psi. Insoluble cell debris was pelleted via centrifugation (30,000 g, 50 min at 4 °C). The resulting supernatant was incubated with 2 ml of a 50% slurry of HisPur Ni-NTA agarose resin (Thermo Fisher Scientific) at 4 °C for 2 hours. The Ni-NTA agarose was pelleted and washed with 30 ml of Ni-NTA wash buffer (50 mM HEPES pH=8], 0.5 M KCl, 1 mM DTT, 25 mM imidazole, and 10% glycerol). Then the DivL-LZ fusion was eluted from the agarose with Ni-NTA elution buffer (50 mM HEPES pH=8.0, 0.5 M KCl, 1 mM DTT, 250 mM imidazole, and 10% glycerol) and concentrated using Amicon Centrifugal Filter Units (30 kDa cutoff), aliquoted and frozen at 80 °C. The concentration was determined using the predicted molecular weight of 58 kDa and 78,630 M-1cm-1.

**Gel Filtration Chromatography:** A gel filtration standard (Sigma) containing thyroglobulin (bovine), apoferritin (horse spleen), β-amylase (sweet potato), alcohol dehydrogenase (yeast), albumin (bovine serum) and carbonic anhydrase (bovine erythrocytes) was used to generate a molecular weight standard plot using a Superdex 200 10-300 GL column (GE Heathcare). A 1.9 mg/mL sample of DivL(1-536)-LZ and a 3.2 mg/mL sample of DivL(1-532)-LZ were loaded onto the column and eluted with 50 mM HEPES pH=8, 0.5 M KCl.

**Table S2. DNA oligos used in this study**

| Name | Description |
|---|---|
| RecXyl-2 | TCTTCCGGCAGGAATTCACTCACGCC |
| RecUni-1 | ATGCCGTTTGTGATGGCTTCCATGTCG |
| WSC10295 | TGGTACCTTAAGATCTCGAGCTATGACTTCGTACGACCTGATCCTCGC |
| WSC10296 | TCGAATTCTCCGGGAAGCCGAGTTCGGGCTGCATGG |
| WSC10297 | TGCATGGTACCTTAAGATCTCGAGCTATGACTTCGTACGACCTGATCCTCGCG |
| WSC10298 | TGACGCGTAACGTTCGAATTCTCCGGCAGCTCCGAATAGCCGATGATCGTCG |
| WSC10299 | TGACGCGTAACGTTCGAATTCTCCGGGGCTTCGGCCAGGGCCGC |
| WSC10303 | TGACGCGTAACGTTCGAATTCTCCGGGGTGACGTCGGCGAAGGCG |
| WSC10304 | CGCGACCCTCGACCCCGTGGGACCACAGGGC |
| WSC10305 | GTGGTCCCACGGGGTCGAGGGTCGCGCGGCG |
| WSC10171 | TGACGCGTAACGTTCGAATTCTCCGGGAAGCCGAGTTCGGGCTGC |
| WSC10301 | CCAGCACGGCTCGACCCCGTGGGACCACAGGGC |
| WSC10302 | GTGGTCCCACGGGGTCGAGCCGTGCTGGATCGC |
| WSC10306 | GGACATCCTCGATCTCCACGCTGTCGACGAAGGCGG |
| WSC10307 | CGTCGACAGCGTGGAGATCGAGGATGTCCGCGACGC |
| WSC10308 | GGCGCAGCTCGCCGGTGACGTCGGCGCAGAACAC |
| WSC10309 | CGCCGACGTCACCGGCGAGCTGCGCCTGAAAGC |
| WSC10310 | GGTCTCGGGTGTCGGTGATGTCGGAATAGATCAGCAGC |
| WSC10311 | TATTCCGACATCACCGACACCCGAGACCTGCAGAGCG |
| KAKqc0018 | GATCGCCTTCGCCGCCGTCACCGACACC |
| KAKqc0019 | GGTGTCGGTGACGGCGGCGAAGGCGATC |
| KAKqc0020 | CTTCGCCGACGTCGTCGACACCCGAGAC |
| KAKqc0021 | GTCTCGGGTGTCGACGACGTCGGCGAAG |
| KAKqc0046 | GGGCTGGTTCGAGGATGCCGCTTAG |
| KAKqc0049 | GGAGAGAGCGAGATTCTCCGCGCTG |
| KAKqc0076 | TGACGCGTAACGTTCGAATTCTCCGGCAGACGGGCGACCTCATTCTCGAGGTGGTAGTT CTTCGAGAGGAGTTCTTCGACCTTGTCTTCGAGTTGCTTGGCTTCGGCCAGGGCCGC |
| KAKqc0077 | TGACGCGTAACGTTCGAATTCTCCGGCAGACGGGCGACCTCATTCTCGAGGTGGTAGTT CTTCGAGAGGAGTTCTTCGACCTTGTCTTCGAGTTGCTTTTCGGCCAGGGCCGCCGAGC |
| KAKqc0078 | TGACGCGTAACGTTCGAATTCTCCGGCAGACGGGCGACCTCATTCTCGAGGTGGTAGTT CTTCGAGAGGAGTTCTTCGACCTTGTCTTCGAGTTGCTTGGCCAGGGCCGCCGAGCG |
| KAKqc0079 | TGACGCGTAACGTTCGAATTCTCCGGCAGACGGGCGACCTCATTCTCGAGGTGGTAGTT CTTCGAGAGGAGTTCTTCGACCTTGTCTTCGAGTTGCTTCAGGGCCGCCGAGCGATC |
| KAKqc0080 | TGACGCGTAACGTTCGAATTCTCCGGCAGACGGGCGACCTCATTCTCGAGGTGGTAGTT CTTCGAGAGGAGTTCTTCGACCTTGTCTTCGAGTTGCTTGGCCGCCGAGCGATCGGC |
| KAKqc0081 | TGACGCGTAACGTTCGAATTCTCCGGCAGACGGGCGACCTCATTCTCGAGGTGGTAGTT CTTCGAGAGGAGTTCTTCGACCTTGTCTTCGAGTTGCTTCGCCGAGCGATCGGCCAGG |
| KAKqc0082 | TGACGCGTAACGTTCGAATTCTCCGGCAGACGGGCGACCTCATTCTCGAGGTGGTAGTT CTTCGAGAGGAGTTCTTCGACCTTGTCTTCGAGTTGCTTCGAGCGATCGGCCAGGGC |
| KAKqc0108 | TGACGCGTAACGTTCGAATTCTCCGGCAGACGGGCGACCTCATTCTCGAGGTGGTAGTT CTTCGAGAGGAGTTCTTCGACCTTGTCTTCGAGTTGCTTCACGCTGTCGACGAAGGC |
| KAK1 | TGCATGGTACCTTAAGATCTCGAGCTATGACTTCGTACGACCTGATCCTCGC |
| KAK2 | TGACGCGTAACGTTCGAATTCTCCGGGAAGCCGAGTTCGGGCTGCATGG |
| KAK3 | CGAGGTCGACGGTATCGATAAGCTTGATATGACTTCGTACGACCTGATCCTCGCG |
| KAK5 | CGGCGCTTTTCCATCGAGAATTCGATGGCTTCGGCCAGGGCCGC |

| | |
|---|---|
| KAK7 | CGGCGCTTTTCCATCGAGAATTCGATGAAGCCGAGTTCGGGCTGCATGG |
| KAK8 | CGGCGCTTTTCCATCGAGAATTCGATGGTGACGTCGGCGAAGGCG |
| KAK9 | ATCGAATTCTCGATGGAAAAGCGCCG |
| KAK10 | ATCAAGCTTATCGATACCGTCGACCTCGAG |
| KAK31 | CGGAAGCAGCTGTCGCACGGCCTCTTAAGCTTGCAATGCGCAGT |
| KAK32 | TGACGCGTAACGTTCGAATTCTCCGGCGAAACAAGGCCATGGGG |
| KAK33 | TGCATGGTACCTTAAGATCTCGAGCTATGCAGCGACGGAACCTGGAGGC |
| KAK34 | GGTCTCGGGTGTCCACGCTGTCGACGAAGGC |
| KAK35 | CGTCGACAGCGTGGACACCCGAGACCTGCAGAGCG |
| KAK36 | GGTCTCGGGTGTCCGAAACAAGGCCATGGGG |
| KAK37 | TGGCCTTGTTTCGGACACCCGAGACCTGCAGAGCG |
| KAK63 | TGACGCGTAACGTTCGAATTCTCCGGGGTGACGTCGGCGAAGGCG |
| KAK64 | TGACGCGTAACGTTCGAATTCTCCGGGGTGATGTCGGAATAGATCAGCAGC |
| KAK91 | TGCATGGTACCTTAAGATCTCGAGCTATGGTCGAGCCGTGCTGGATCGC |
| KAK92 | TGACGCGTAACGTTCGAATTCTCCGGGGTGATGTCGGAATAGATCAGCAGC |
| KAK93 | TGCATGGTACCTTAAGATCTCGAGCTATGGAGATCGAGGATGTCCGCGACGC |
| KAK94 | TGACGCGTAACGTTCGAATTCTCCGGGGTGACGTCGGCGAAGGCG |
| KAK132 | CGGCGCTTTTCCATCGAGAATTCGATCAGACGGGCGACCTCATTCTCGAGG |
| KAK155 | CCAACTAGTGAAAACCTGTATTTTCAGGGCGCTATGGCCTGGCTCGACGCCTTCGA |
| KAK158 | GCTCGAGAATTCCATGGCCATATGGCTTCACAGACGGGCGACCTCATTCTCGA |
| KAK173 | GCCCGAACTCGGCTTCTTAAGATCTCGAGCTCCGGAGAATTCG |
| KAK174 | GGTCGTACGAAGTCATGGTACCATGCATATTAATTAAGGCGCCTGC |
| KAK175 | CCTTAATTAATATGCATGGTACCATGACTTCGTACGACCTGATCCTCGC |
| KAK176 | GCTCGAGATCTTAAGAAGCCGAGTTCGGGCTGCATGG |
| KAK177 | CGAACTCGGCTTCCACCGGTCGGCCACCATGG |
| KAK178 | GGTCGTACGAAGTCATACGCGTAACGTTCGAATTCTCCGG |
| KAK179 | TCGAACGTTACGCGTATGACTTCGTACGACCTGATCCTCGC |
| KAK180 | TGGCCGACCGGTGGAAGCCGAGTTCGGGCTGCATGG |
| KAK181 | CGGCTTCGGATCCATGGTGAGCAAGGGCGAGGAGG |
| KAK182 | CGTACGAAGTCATGGTGGCCGACCGGTGACG |
| KAK183 | CCGGTCGGCCACCATGACTTCGTACGACCTGATCCTCGC |
| KAK184 | CCTTGCTCACCATGGATCCGAAGCCGAGTTCGGGCTGCATGG |
| KAK189 | ACTCGGCTTCTAACCTGCAGGCGCCTTAATTAATATGCATGG |
| KAK190 | TGGCCGACCGGTGCTTGTACAGCTCGTCCATGCCGCC |
| KAK191 | CGAGCTGTACAAGCACCGGTCGGCCACCATGACTTCGTACGACCTGATCCTCGC |
| KAK192 | AGGCGCCTGCAGGTTAGAAGCCGAGTTCGGGCTGCATGG |
| KAK323 | TTTTTGGGCTAACAGGAGGAATTAACCAATGGTCGAGGGTCGCGCGGCG |
| KAK324 | TTACTGCCGCCGCCGCCGCTCTCGAGGAAGCCGAGTTCGGGCTGCATGG |
| KAK325 | TTTTTGGGCTAACAGGAGGAATTAACCAATGGAGGCGCGTATCGTCGCGC |
| KAK326 | TTACTGCCGCCGCCGCCGCTCTCGAGCGAAACAAGGCCATGGGGACCG |

**Table S3. Gibson cloning strategy to generate plasmids expressing DivL variants**

| Plasmid | Plasmid Description | First Insert Forward Primer | First Insert Reverse Primer | Second Insert Forward Primer | Second Insert Reverse Primer |
|---|---|---|---|---|---|
| WSC10153 | pXCHYC-6 DivL (1-769) | WSC10295 | WSC10296 | | |
| pWSC10155 | pXCHYC-6 DivL (1-537) | WSC10297 | WSC10299 | | |
| pWSC10157 | pXCHYC-6 DivL (1-27, 171-769) | WSC10297 | WSC10301 | WSC10302 | WSC10171 |
| pWSC10158 | pXCHYC-6 DivL (1-518) | WSC10297 | WSC10303 | | |
| pWSC10159 | pXCHYC-6 DivL (1-27, 134-769) | WSC10297 | WSC10304 | WSC10305 | WSC10171 |
| pWSC10160 | pXCHYC-6 DivL (1-170, 264-769) | WSC10297 | WSC10306 | WSC10307 | WSC10171 |
| pWSC10161 | pXCHYC-6 DivL (1-262, 392-769) | WSC10297 | WSC10308 | WSC10309 | WSC10171 |
| pWSC10162 | pXCHYC-6 DivL (1-391, 519-769) | WSC10297 | WSC10310 | WSC10311 | WSC10171 |
| pKAK0009 | pXCHYC-6 DivL D516A | WSC10295 | KAKqc0019 | KAKqc0018 | WSC10296 |
| pKAK0010 | pXCHYC-6 DivL T518V | WSC10295 | KAKqc0021 | KAKqc0020 | WSC10296 |
| pKAK0012 | pBXMCS-2 DivL (1-537) | KAK3 | KAK5 | | |
| pKAK0015 | pBXMCS-2 DivL (1-518) | KAK3 | KAK8 | | |
| pKAK0016 | pBXMCS-2 DivL (1-27, 134-769) | KAK3 | KAK7 | | |
| pKAK0017 | pBXMCS-2 DivL (1-170, 264-769) | KAK3 | KAK7 | | |
| pKAK0018 | pBXMCS-2 DivL (1-262, 392-769) | KAK3 | KAK7 | | |
| pKAK0019 | pBXMCS-2 DivL (1-391, 519-769) | KAK3 | KAK7 | | |
| pKAK0037 | pBXMCS-2 DivL D516A | KAK3 | KAK7 | | |
| pKAK0038 | pBXMCS-2 DivL T518V | KAK3 | KAK7 | | |
| pKAK0045 | pXCHYC-6 DivL (1-170) | KAK1 | KAK31 | | |
| pKAK0046 | pXCHYC-6 DivL (1-133) | KAK1 | KAK32 | | |
| pKAK0047 | pXCHYC-6 DivL (28-170) | KAK33 | KAK31 | | |
| pKAK0048 | pXCHYC-6 DivL (28-133) | KAK33 | KAK32 | | |
| pKAK0049 | pXCHYC-6 DivL (1-170, 519-769) | KAK1 | KAK34 | KAK35 | KAK2 |
| pKAK0050 | pXCHYC-6 DivL (1-133, 519-769) | KAK1 | KAK36 | KAK37 | KAK2 |
| pKAK0051 | pXCHYC-6 DivL (28-170, 519-769) | KAK33 | KAK34 | KAK35 | KAK2 |
| KAK0052 | pXCHYC-6 DivL (28-170, 519-769) | KAK33 | KAK36 | KAK37 | KAK2 |
| pKAK0072 | pXCHYC-6 DivL (1-170, 263-518) | KAK1 | WSC10306 | WSC10307 | KAK63 |
| pKAK0073 | pXCHYC-6 DivL (1-391) | KAK1 | KAK64 | | |

| | | | |
|---|---|---|---|
| pKAK0079 | pXCHYC-6 DivL (1-537, LZ) | KAK1 | KAKqc0076 |
| pKAK0080 | pXCHYC-6 DivL (1-536, LZ) | KAK1 | KAKqc0077 |
| pKAK0081 | pXCHYC-6 DivL (1-535, LZ) | KAK1 | KAKqc0078 |
| pKAK0082 | pXCHYC-6 DivL (1-534, LZ) | KAK1 | KAKqc0079 |
| pKAK0083 | pXCHYC-6 DivL (1-533, LZ) | KAK1 | KAKqc0080 |
| pKAK0084 | pXCHYC-6 DivL (1-532, LZ) | KAK1 | KAKqc0081 |
| pKAK0085 | pXCHYC-6 DivL (1-531, LZ) | KAK1 | KAKqc0082 |
| pKAK0088 | pXCHYC-6 DivL (171-391) | KAK91 | KAK92 |
| pKAK0089 | pXCHYC-6 DivL (263-518) | KAK93 | KAK94 |
| pKAK0096 | pXCHYC-6 DivL (1-170, LZ) | KAK1 | KAKqc0108 |
| pKAK0121a | pBXMCS-2 DivL (1-537, LZ) | KAK3 | KAK132 |
| pKAK0121b | pBXMCS-2 DivL (1-536, LZ) | KAK3 | KAK132 |
| pKAK0121c | pBXMCS-2 DivL (1-535, LZ) | KAK3 | KAK132 |
| pKAK0121d | pBXMCS-2 DivL (1-534, LZ) | KAK3 | KAK132 |
| pKAK0121e | pBXMCS-2 DivL (1-533, LZ) | KAK3 | KAK132 |
| pKAK0121f | pBXMCS-2 DivL (1-532, LZ) | KAK3 | KAK132 |
| pKAK0121g | pBXMCS-2 DivL (1-531, LZ) | KAK3 | KAK132 |
| pKAK0137b | pTEV-5 DivL (1-532, LZ) | KAK155 | KAK158 |
| pKAK0137f | pTEV-5 DivL (1-536, LZ) | KAK155 | KAK158 |
| pKAK0141 | pXCHYC-6 DivL LRSRAPENSNVT RHRSAT mCherry linker | KAK175 | KAK176 |
| pKAK0142 | pXCHYC-6 DivL HRSAT mCherry linker | KAK179 | KAK180 |
| pKAK0143 | pXCHYC-6 DivL GS mCherry linker | KAK183 | KAK184 |
| pKAK0145 | pXCHYN-6 DivL mCherry HRSAT linker | KAK191 | KAK192 |
| pKAK0198 | pXCHYC-6 DivL (1-532, LZ) D516A | KAK1 | KAKqc0081 |
| pKAK0199 | pXCHYC-6 DivL (1-532, LZ) T518V | KAK1 | KAKqc0081 |
| pKAK0207 | pBAD DivL (134-769), msfGFP | KAK323 | KAK234 |
| pKAK0208 | pBAD DivL (33-133), msfGFP | KAK325 | KAK326 |

**Table S4. Plasmids used in this study**

| Plasmid | Description | Reference |
|---|---|---|
| pET-28b(+) | bacterial expression vector | Novagen |
| pTEV-5 | bacterial expression vector | (4) |
| pXCHYC-6 | C. crescentus integrating C-terminal mCherry fusion vector | (5) |
| pBXMCS-2 | C. crescentus high-copy replicating C-terminal M2 fusion vector | (5) |
| pWZ013-30 | pCDFDuet1-mcherry-PodJ | Unpublished W. Zhao, S. Childers |
| WSC10153 | pXCHYC-6 DivL (1-769) | This study |
| pWSC10155 | pXCHYC-6 DivL (1-537) | This study |
| pWSC10157 | pXCHYC-6 DivL (1-27, 171-769) | This study |
| pWSC10158 | pXCHYC-6 DivL (1-518) | This study |
| pWSC10159 | pXCHYC-6 DivL (1-27, 134-769) | This study |
| pWSC10160 | pXCHYC-6 DivL (1-170, 264-769) | This study |
| pWSC10161 | pXCHYC-6 DivL (1-262, 392-769) | This study |
| pWSC10162 | pXCHYC-6 DivL (1-391, 519-769) | This study |
| pKAK0009 | pXCHYC-6 DivL D516A | This study |
| pKAK0010 | pXCHYC-6 DivL T518V | This study |
| pKAK0012 | pBXMCS-2 DivL (1-537) | This study |
| pKAK0015 | pBXMCS-2 DivL (1-518) | This study |
| pKAK0016 | pBXMCS-2 DivL (1-27, 134-769) | This study |
| pKAK0017 | pBXMCS-2 DivL (1-170, 264-769) | This study |
| pKAK0018 | pBXMCS-2 DivL (1-262, 392-769) | This study |
| pKAK0019 | pBXMCS-2 DivL (1-391, 519-769) | This study |
| pKAK0037 | pBXMCS-2 DivL D516A | This study |
| pKAK0038 | pBXMCS-2 DivL T518V | This study |
| pKAK0045 | pXCHYC-6 DivL (1-170) | This study |
| pKAK0046 | pXCHYC-6 DivL (1-133) | This study |
| pKAK0047 | pXCHYC-6 DivL (28-170) | This study |
| pKAK0048 | pXCHYC-6 DivL (28-133) | This study |
| pKAK0049 | pXCHYC-6 DivL (1-170, 519-769) | This study |
| pKAK0050 | pXCHYC-6 DivL (1-133, 519-769) | This study |
| pKAK0051 | pXCHYC-6 DivL (28-170, 519-769) | This study |
| pKAK0052 | pXCHYC-6 DivL (28-133, 519-769) | This study |
| pKAK0072 | pXCHYC-6 DivL (1-170, 263-518) | This study |
| pKAK0073 | pXCHYC-6 DivL (1-391) | This study |
| pKAK0079 | pXCHYC-6 DivL (1-537, LZ) | This study |
| pKAK0080 | pXCHYC-6 DivL (1-536, LZ) | This study |
| pKAK0081 | pXCHYC-6 DivL (1-535, LZ) | This study |
| pKAK0082 | pXCHYC-6 DivL (1-534, LZ) | This study |

| | | |
|---|---|---|
| pKAK0083 | pXCHYC-6 DivL (1-533, LZ) | This study |
| pKAK0084 | pXCHYC-6 DivL (1-532, LZ) | This study |
| pKAK0085 | pXCHYC-6 DivL (1-531, LZ) | This study |
| pKAK0088 | pXCHYC-6 DivL (171-391) | This study |
| pKAK0089 | pXCHYC-6 DivL (263-518) | This study |
| pKAK0096 | pXCHYC-6 DivL (1-170, LZ) | This study |
| pKAK0121a | pBXMCS-2 DivL (1-537, LZ) | This study |
| pKAK0121b | pBXMCS-2 DivL (1-536, LZ) | This study |
| pKAK0121c | pBXMCS-2 DivL (1-535, LZ) | This study |
| pKAK0121d | pBXMCS-2 DivL (1-534, LZ) | This study |
| pKAK0121e | pBXMCS-2 DivL (1-533, LZ) | This study |
| pKAK0121f | pBXMCS-2 DivL (1-532, LZ) | This study |
| pKAK0121g | pBXMCS-2 DivL (1-531, LZ) | This study |
| pKAK0137b | pTEV-5 DivL (54-532, LZ) | This study |
| pKAK0137f | pTEV-5 DivL (54-536, LZ) | This study |
| pKAK0141 | pXCHYC-6 DivL LRSRAPENSNVTRHRSAT mCherry linker | This study |
| pKAK0142 | pXCHYC-6 DivL HRSAT mCherry linker | This study |
| pKAK0143 | pXCHYC-6 DivL GS mCherry linker | This study |
| pKAK0145 | pXCHYN-6 DivL mCherry HRSAT linker | This study |
| pKAK0198 | pXCHYC-6 DivL (1-532, LZ) D516A | This study |
| pKAK0199 | pXCHYC-6 DivL (1-532, LZ) T518V | This study |
| pKAK0207 | pBAD DivL (134-769), msfGFP | This study |
| pKAK0208 | pBAD DivL (33-133), msfGFP | This study |

**Table S5. *C. crescentus* and *E. coli* strains used in this study**

| Strain | Description | Plasmid(s) | Reference Source |
|---|---|---|---|
| *E. coli* DH5α | bacterial cloning strain | | Invitrogen |
| *E. coli* BL21 | bacterial expression strain | | Novagen |
| *C. crescentus* NA1000 | laboratory Caulobacter crescentus strain | | Shapiro Lab |
| WSC0427 | Δ*divL*(strep) | pMR20-divL | (6) Shapiro Lab |
| WSC1131 | *DivK341* DivK$^{cs}$ | | (7) Shapiro Lab |
| JH42 | BL21(DE3) pACYC-mCherry-PopZ | | (8) Bowman Lab |
| JH79 | BL21 (DE3) pACYC-mCherry-PopZ + pBAD-DivL-msfGFP | | (8) Bowman Lab |
| WSC0300 | C. crescentus pBXMCS-2 | | This study |
| WSC0302 | *C. crescentus* pBXMCS-2 DivL (1-769) | | This study |
| WSC1233 | DH5α pCDFDuet1-mcherry-PodJ | pWZ013-30 | Unpublished W. Zhao, S. Childers |
| WSC1368 | BL21(DE3) pCDFDuet1-mcherry-PodJ | pWZ013-30 | Unpublished W. Zhao, S. Childers |
| WSC0454 | *C. crescentus* pXCHYC-6 DivL (1-537) | pWSC10155 | This study |
| WSC0457 | *C. crescentus* pXCHYC-6 DivL (1-518) | pWSC10158 | This study |
| WSC0458 | *C. crescentus* pXCHYC-6 DivL (1-27, 134-769) | pWSC10159 | This study |
| WSC0459 | *C. crescentus* pXCHYC-6 DivL (1-170, 264-769) | pWSC10160 | This study |
| WSC0460 | *C. crescentus* pXCHYC-6 DivL (1-262, 392-769) | pWSC10161 | This study |
| WSC0461 | *C. crescentus* pXCHYC-6 DivL (1-391, 519-769) | pWSC10162 | This study |
| KAK12 | DH5α pXCHYC-6 DivL (1-537) | pWSC10155 | This study |
| KAK13 | DH5α pXCHYC-6 DivL (1-27, 171-769) | pWSC10157 | This study |
| KAK14 | DH5α pXCHYC-6 DivL (1-518) | pWSC10158 | This study |
| KAK15 | DH5α pXCHYC-6 DivL (1-27, 134-769) | pWSC10159 | This study |
| KAK16 | DH5α pXCHYC-6 DivL (1-170, 264-769) | pWSC10160 | This study |
| KAK17 | DH5α pXCHYC-6 DivL (1-262, 392-769) | pWSC10161 | This study |
| KAK18 | DH5α pXCHYC-6 DivL (1-391, 519-769) | pWSC10162 | This study |
| KAK20 | DH5α pXCHYC-6 DivL (1-769) | pWSC10153 | This study |
| KAK30 | DH5α pBXMCS-2 DivL D516A | pKAK0037 | This study |
| KAK33 | DH5α pBXMCS-2 DivL T518V | pKAK0038 | This study |
| KAK34 | DH5α pXCHYC-6 DivL D516A | pKAK0009 | This study |
| KAK36 | DH5α pBXMCS-2 DivK | pBXSPA-DivK | This study |
| KAK43 | NA1000 pXCHYC-6 DivL (1-769) | WSC10153 | This study |
| KAK45 | DH5α pBXMCS-2 DivL (1-537) | pKAK0012 | This study |
| KAK47 | DH5α pBXMCS-2 DivL (1-518) | pKAK0015 | This study |
| KAK48 | DH5α pBXMCS-2 DivL (1-27, 134-769) | pKAK0016 | This study |
| KAK49 | DH5α pBXMCS-2 DivL (1-391, 519-769) | pKAK0019 | This study |
| KAK51 | DH5α pBXMCS-2 DivL (1-170, 264-769) | pKAK0017 | This study |
| KAK52 | DH5α pBXMCS-2 DivL (1-262, 392-769) | pKAK0018 | This study |
| KAK62 | NA1000 pXCHYC-6 DivL HRSAT mCherry linker | pKAK0142 | This study |

| | | | |
|---|---|---|---|
| KAK63 | DH5α pXCHYC-6 DivL T518V | pKAK0010 | This study |
| KAK64 | NA1000 pBXMCS-2 DivL (1-537) | pKAK00012 | This study |
| KAK65 | NA1000 pBXMCS-2 DivL (1-518) | pKAK0015 | This study |
| KAK66 | NA1000 pBXMCS-2 DivL (1-262, 392-769) | pKAK0018 | This study |
| KAK67 | NA1000 pBXMCS-2 DivL (1-391, 519-769) | pKAK0019 | This study |
| KAK68 | NA1000 pBXMCS-2 DivL (1-27, 134-769) | pKAK0016 | This study |
| KAK69 | NA1000 pBXMCS-2 DivL (1-170, 264-769) | pKAK0017 | This study |
| KAK92 | NA1000 pXCHYC-6 DivL D516A | pKAK0009 | This study |
| KAK117 | DH5α pXCHYC-6 DivL (28-170) | pKAK0047 | This study |
| KAK102 | NA1000 pXCHYC-6 DivL T518V | pKAK0010 | This study |
| KAK106 | NA1000 pXCHYC-6 DivL (1-27, 171-769) | pWSC10157 | This study |
| KAK115 | NA1000 Δ*divL* pXCHYC-6 DivL (1-769) | pWSC10153 | This study |
| KAK119 | DH5α pXCHYC-6 DivL (28-133) | pKAK0048 | This study |
| KAK122 | DH5α pXCHYC-6 DivL (1-170) | pKAK0045 | This study |
| KAK123 | DH5α pXCHYC-6 DivL (1-133) | pKAK0046 | This study |
| KAK124 | DH5α pXCHYC-6 DivL (1-170, 519-769) | pKAK0049 | This study |
| KAK125 | DH5α pXCHYC-6 DivL (1-133, 519-769) | pKAK0050 | This study |
| KAK126 | DH5α pXCHYC-6 DivL (28-170, 519-769) | pKAK0051 | This study |
| KAK127 | DH5α pXCHYC-6 DivL (28-133, 519-769) | pKAK0052 | This study |
| KAK140 | NA1000 pXCHYC-6 DivL (1-170) | pKAK0045 | This study |
| KAK141 | NA1000 pXCHYC-6 DivL (1-133) | pKAK0046 | This study |
| KAK142 | NA1000 pXCHYC-6 DivL (28-170) | pKAK0047 | This study |
| KAK143 | NA1000 pXCHYC-6 DivL (28-133) | pKAK0048 | This study |
| KAK144 | NA1000 pXCHYC-6 DivL (1-170, 519-769) | pKAK0049 | This study |
| KAK145 | NA1000 pXCHYC-6 DivL (1-133, 519-769) | pKAK0050 | This study |
| KAK146 | NA1000 pXCHYC-6 DivL (28-170, 519-769) | pKAK0051 | This study |
| KAK147 | NA1000 pXCHYC-6 DivL (28-133, 519-769) | pKAK0052 | This study |
| KAK150 | NA1000 pBXMCS-2 DivL D516A | pKAK0037 | This study |
| KAK151 | NA1000 pBXMCS-2 DivL T518V | pKAK0038 | This study |
| KAK172 | DH5α pXCHYC-6 DivL (1-537, LZ) | pKAK0079 | This study |
| KAK176 | DH5α pXCHYC-6 DivL (1-536, LZ) | pKAK0080 | This study |
| KAK177 | DH5α pXCHYC-6 DivL (1-535, LZ) | pKAK0081 | This study |
| KAK178 | DH5α pXCHYC-6 DivL (1-534, LZ) | pKAK0082 | This study |
| KAK179 | DH5α pXCHYC-6 DivL (1-533, LZ) | pKAK0083 | This study |
| KAK180 | DH5α pXCHYC-6 DivL (1-532, LZ) | pKAK0084 | This study |
| KAK181 | DH5α pXCHYC-6 DivL (1-531, LZ) | pKAK0085 | This study |
| KAK186 | DH5α pXCHYC-6 DivL (1-170, 263-518) | pKAK0072 | This study |
| KAK188 | DH5α pXCHYC-6 DivL (171-391) | pKAK0088 | This study |
| KAK191 | DH5α pXCHYC-6 DivL (171-391) | pKAK0089 | This study |
| KAK197 | NA1000 pXCHYC-6 DivL (1-536, LZ) | pKAK0080 | This study |
| KAK198 | NA1000 pXCHYC-6 DivL (1-535, LZ) | pKAK0081 | This study |
| KAK199 | NA1000 pXCHYC-6 DivL (1-533, LZ) | pKAK0083 | This study |

| | | | |
|---|---|---|---|
| KAK200 | NA1000 pXCHYC-6 DivL (1-532, LZ) | pKAK0084 | This study |
| KAK201 | NA1000 pXCHYC-6 DivL (1-531, LZ) | pKAK0085 | This study |
| KAK202 | NA1000 pXCHYC-6 DivL (1-537, LZ) | pKAK0079 | This study |
| KAK205 | NA1000 pXCHYC-6 DivL (1-534, LZ) | pKAK0082 | This study |
| KAK209 | DH5α pBXMCS-2 DivL (1-537, LZ) | pKAK0121a | This study |
| KAK214 | DH5α pBXMCS-2 DivL (1-536, LZ) | pKAK0121b | This study |
| KAK215 | NA1000 pXCHYC-6 DivL (1-170, 263-518) | pKAK0072 | This study |
| KAK221 | NA1000 pXCHYC-6 DivL (171-391) | pKAK0088 | This study |
| KAK222 | NA1000 pXCHYC-6 DivL (263-518) | pKAK0089 | This study |
| KAK228 | NA1000 pXCHYC-6 DivL (1-391) | pKAK0073 | This study |
| KAK229 | DH5α pBXMCS-2 DivL (1-535, LZ) | pKAK0121c | This study |
| KAK230 | DH5α pBXMCS-2 DivL (1-531, LZ) | pKAK0121g | This study |
| KAK232 | DH5α pXCHYN-6 DivL mCherry HRSAT linker | pKAK0145 | This study |
| KAK254 | DH5α pTEV-5 DivL (54-536, LZ) | pKAK0137b | This study |
| KAK266 | DH5α pXCHYC-6 DivL LRSRAPENSNVTRHRSAT mCherry linker | pKAK0141 | This study |
| KAK267 | DH5α pXCHYC-6 DivL GS mCherry linker | pKAK0143 | This study |
| KAK270 | DH5α pTEV-5 DivL (54-532, LZ) | pKAK0137f | This study |
| KAK276 | DH5α pBXMCS-2 DivL (1-534, LZ) | pKAK0121d | This study |
| KAK277 | DH5α pXCHYC-6 DivL HRSAT mCherry linker | pKAK0142 | This study |
| KAK281 | NA1000 pXCHYC-6 DivL LRSRAPENSNVTRHRSAT mCherry linker | pKAK0141 | This study |
| KAK283 | NA1000 pXCHYC-6 DivL GS mCherry linker | pKAK0143 | This study |
| KAK293 | DH5α pBXMCS-2 DivL (1-533, LZ) | pKAK0121e | This study |
| KAK294 | DH5α pBXMCS-2 DivL (1-532, LZ) | pKAK0121f | This study |
| KAK299 | NA1000 Δ*divL* pXCHYC-6 DivL HRSAT mCherry linker | pKAK0142 | This study |
| KAK300 | NA1000 Δ*divL* pXCHYC-6 DivL GS mCherry linker | pKAK0143 | This study |
| KAK303 | DH5α pXCHYC-6 DivL (1-391) | pKAK0073 | This study |
| KAK309 | NA1000 pBXMCS-2 DivL (1-536, LZ) | pKAK0121b | This study |
| KAK310 | NA1000 pBXMCS-2 DivL (1-535, LZ) | pKAK0121c | This study |
| KAK311 | NA1000 pBXMCS-2 DivL (1-533, LZ) | pKAK0121e | This study |
| KAK312 | NA1000 pBXMCS-2 DivL (1-532, LZ) | pKAK0121f | This study |
| KAK313 | NA1000 pBXMCS-2 DivL (1-531, LZ) | pKAK0121g | This study |
| KAK324 | NA1000 pBXMCS-2 DivL (1-537, LZ) | pKAK0121a | This study |
| KAK332 | BL21 pTEV-5 DivL (54-536, LZ) | pKAK0137b | This study |
| KAK334 | BL21 pTEV-5 DivL (54-532, LZ) | pKAK0137f | This study |
| KAK336 | NA1000 pBXMCS-2 DivL (1-534, LZ) | pKAK0121d | This study |
| KAK340 | NA1000 pXCHYN-6 DivL mCherry HRSAT linker | pKAK0145 | This study |
| KAK351 | DH5α pXCHYC-6 DivL (1-170, LZ) | pKAK0096 | This study |
| KAK356 | DH5α pXCHYC-6 DivL (1-532, LZ) D516A | pKAK0198 | This study |
| KAK357 | DH5α pXCHYC-6 DivL (1-532, LZ) T518V | pKAK0199 | This study |
| KAK358 | NA1000 pXCHYC-6 DivL (1-170, LZ) | pKAK0096 | This study |

| | | | |
|---|---|---|---|
| KAK363 | NA1000 pXCHYC-6 DivL (1-532, LZ) D516A | pKAK0198 | This study |
| KAK364 | NA1000 pXCHYC-6 DivL (1-532, LZ) T518V | pKAK0199 | This study |
| KAK392 | DH5α pBAD DivL (134-769), msfGFP | pKAK0207 | This study |
| KAK393 | DH5α pBAD DivL (33-133), msfGFP | pKAK0208 | This study |
| KAK406 | BL21 pBAD DivL (33-133), msfGFP + pACYC-mCherry-PopZ | pKAK0208, pACYC-mCherry-PopZ | This study |
| KAK411 | BL21 pBAD DivL (33-133), msfGFP | pKAK0208 | This study |
| KAK423 | BL21 pBAD DivL (134-769), msfGFP +pCDFDuet1-mcherry-PodJ | pKAK0207, pWZ013-30 | This study |
| KAK424 | BL21 pBAD DivL (33-133), msfGFP +pCDFDuet1-mcherry-PodJ | pKAK0208, pWZ013-30 | This study |
| KAK426 | BL21 pBAD DivL (1-769), msfGFP | pBAD-DivL-WT-msfGFP, pWZ013-30 | This study |
| KAK427 | BL21 pBAD DivL (134-769), msfGFP | pKAK0207 | This study |
| KAK428 | BL21 pBAD DivL (134-769), msfGFP +pACYC-mCherry-PopZ | pKAK0207, pACYC-mCherry-PopZ | This study |
| KAK460 | BL21 pBAD DivL, msfGFP | pBAD-DivL-msfGFP | This study |
| KAK463 | DH5α pBAD DivL, msfGFP | pBAD-DivL-msfGFP | This study |
